# Spatial metabolomics links polyamine turnover and vascular lipid remodeling to barley spikelet fate

**DOI:** 10.64898/2026.09.23.753928

**Authors:** Nandhakumar Shanmugaraj, Yongyu Huang, Adriana Garibay-Hernández, Twan Rutten, John C. D’Auria, Hans-Peter Mock, Thorsten Schnurbusch

## Abstract

Local metabolic programs can determine whether developing organs maintain growth, differentiate, or degenerate. However, the spatial establishment of such programs during reproductive development remains poorly understood. In cereal inflorescences, this question is closely tied to grain number because the fate of initiated floral primordia is shaped by the developmental position, metabolic support, genotype, and environmental stress. To understand the metabolic logic underlying floral fate in barley (*Hordeum vulgare* L.), we mapped the spatial distribution of metabolites across the inflorescence in wild-type Bowman and in *hvcmf4*, a mutant that undergoes premature apical inflorescence degeneration. Amino acid-, carbohydrate-, and chlorophyll-associated metabolites form clear developmental gradients along the inflorescence axis. These gradients are progressively remodeled during developmental pre-anthesis tip degeneration but are both accelerated and spatially distorted in *hvcmf4*. Within this broader metabolic landscape, two discrete domains stand out. First, spermidine-associated domains specifically marked reproductive meristems, a pattern corroborated by the meristem-enriched expression of polyamine biosynthetic genes. This polyamine-rich meristem state declined as developmental tip degeneration proceeded and was prematurely lost in *hvcmf4*, where polyamine metabolism appeared to shift toward oxidative breakdown via the polyamine oxidase pathway. Second, lysophosphatidylcholine accumulated specifically at floral attachment and vascular supply zones. The developmental reduction of this bioactive lysophospholipid-associated domain in the apical regions, and its stronger disruption in *hvcmf4*, coincided with the reprogramming of lipid-remodeling and auxin transport-associated genes, suggesting impaired coordination of lipid signaling, auxin canalization, and vascular support. Together, our study reveals spatial polyamine and lysophospholipid domains that couple meristem maintenance with vascular support, providing a metabolic framework for the fate of developing cereal inflorescences.

## Introduction

Plant organ development depends on local molecular environments that determine whether cells remain meristematic, initiate organ primordia, differentiate, or enter degeneration (Lindsay et al., 2024). Although genetic and transcriptomic approaches have identified many key regulators of this process, the spatial organization of metabolism remains poorly understood (Barmukh et al., 2025). This gap is significant because metabolites are not merely substrates and products of cellular activity; they also function as signaling molecules, redox buffers, membrane-remodeling factors, and regulators of cellular viability (Cai et al., 2023).

Polyamines, including putrescine, spermidine, and spermine, are abundant polycationic metabolites with broad roles in plant development, stress adaptation, and cellular homeostasis (Blázquez, 2024; Yang et al., 2024). Owing to their positive charge, they readily interact with negatively charged macromolecules, DNA, RNA, ribosomes, proteins, phospholipids, and membranes influencing translation, membrane stability, ion channel activity, autophagy, senescence, redox balance, and stress responses. Polyamine oxidases can further couple spermidine and spermine catabolism to hydrogen peroxide production and redox signaling, linking polyamine turnover to cellular viability (Blázquez, 2024; Blagojević et al., 2026). Altered polyamine metabolism has been associated with stress resilience, grain filling, grain quality, and crop productivity, whereas polyamine imbalance can impair fertility or promote cell death depending on the dose, tissue, and developmental context (Burke et al., 2024; Yang et al., 2024; Blagojević et al., 2026). Lipid signaling has gained increasing attention as a complementary regulatory layer that connects membrane dynamics with hormone responses, stress adaptation, and organ growth (Hoffmann-Benning, 2020; Hernández-Esquivel et al., 2025). Central to this regulation is membrane lipid remodeling, defined as the enzymatic turnover and reassembly of phospholipids including phospholipase-mediated deacylation of phosphatidylcholine to generate lysophospholipids, such as lysophosphatidylcholine (LPC), and LPCAT/LPLAT-mediated reacylation, through the Lands cycle to restore phosphatidylcholine homeostasis (Bates et al., 2007; Chang et al., 2026). Therefore, LPC accumulation can indicate local phospholipid turnover, membrane remodeling, or lipid signaling activity (Drissner et al., 2007; Labusch et al., 2013; Poulsen et al., 2015). Despite the established roles of both polyamines and LPC in plant physiology, the spatial positioning of these metabolic states within young cereal reproductive tissues and their shifts during spikelet meristem maintenance or degeneration remain unknown.

Conventional bulk metabolite profiling averages signals across neighboring tissues with distinct developmental fates, obscuring the local metabolic states that may directly influence organogenesis. Recent advances in single-cell and spatial omics have highlighted the importance of resolving molecular programs in their anatomical context, particularly in crops in which organ architecture and developmental position strongly influence yield-related traits (Barmukh et al., 2025). In cereals, single-cell and spatial transcriptome resources have begun to define cell types, meristem subdomains, vascular territories, and regional gene expression programs during inflorescence development (Xu et al., 2025a; Xu et al., 2025b; Demesa-Arevalo et al., 2026; Long et al., 2026). However, gene expression alone does not capture the metabolic states that directly reflect energy availability, redox balance, lipid signaling, and cellular physiology. Spatial metabolomics fills this gap by revealing where metabolites accumulate, decline, or are remodeled within developing organs and by indicating which pathways are most tractable to manipulate (Alexandrov, 2020; Liu et al., 2026). Matrix-assisted laser desorption/ionization mass spectrometry imaging (MALDI-MSI) enables metabolite and lipid visualization directly in tissue sections while preserving the full anatomical context (Kompauer et al., 2017) and has been applied in plants to reveal spatial heterogeneity in seeds, roots, leaves, nodules, and reproductive tissues (Liu et al., 2026). Importantly, recent work has moved beyond descriptive metabolite mapping toward the use of chemical imaging as a tool for developmental hypothesis generation. DESI-MSI of developing maize roots resolved small-molecule gradients across the stem cell-to-differentiation axis and demonstrated that spatially patterned TCA-cycle metabolites influence root meristem activity and differentiation (Zhang et al., 2023), while MSI identified itaconate as an endogenous plant metabolite with roles in growth, oxidative stress, and hormone signaling (Zhang et al., 2025). Despite this progress, the application of MSI to early cereal inflorescence meristems remain limited, owing to their anatomical complexity and rapid developmental transitions. Integrating MALDI-MSI with genetic, transcriptomic, and spatial transcriptome resources therefore offers a direct path toward connecting regulatory domains with the metabolic states that sustain or undermine reproductive organ maintenance and degeneration.

Cereal inflorescences provide a powerful and agronomically relevant system for studying the balance between organ initiation and organ loss during reproductive development (Kellogg et al., 2013). Recent studies on cereal floral development have emphasized that reproductive success depends not only on the initiation of floral organs but also on their maintenance, with vascular patterning and chloroplast maturation emerging as key components of floral fate (Huang and Schnurbusch, 2024). Among cereals, the *Triticeae*, including barley, wheat, and rye, characteristically bear spike-type inflorescences in which spikelets, the basic grain-bearing floral units, are arranged in an alternate-distichous pattern directly along the central rachis axis. Barley exemplifies this organization: its indeterminate spike produces three spikelets (one central and two lateral) at each rachis node, each uniflorous and capable of yielding a single grain when fertile (Koppolu and Schnurbusch, 2019). In barley spikes, a proportion of apical spikelets naturally undergo pre-anthesis tip degeneration (PTD), reducing the final spike fertility and grain number (Shanmugaraj et al., 2023b). This creates a built-in developmental gradient within a single organ; basal and central spikelets remain viable, whereas apical spikelets progressively lose their growth potential. A similar spatial logic is emerging in wheat, where spikelet abortion is accompanied by spatially heterogeneous metabolic reprogramming across the basal, central, and apical spike regions (Dai et al., 2026). These findings suggest that cereal spikelet fate is strongly shaped by the local metabolic state, yet the specific metabolite domains associated with meristem viability and vascular support remain poorly resolved.

Recent genetic, anatomical, and multi-omic studies have shown that spikelet initiation and apical spikelet survival are partially separable processes in barley plants. The CCT motif family gene *HvCMF4*, identified through the premature apical spike degeneration mutant *hvcmf4*, acts after early spikelet initiation to promote spikelet growth, rachis greening, vascular development, carbohydrate metabolism, hormone homeostasis, and spikelet survival (Huang et al., 2023).

HvCMF4 loss caused premature and enhanced spike PTD, reduced rachis greening, altered sugar metabolism, and impaired vascular development. Anatomical analyses have further shown that spike fertility correlates more strongly with vascular area and vein dimensions than with vein number, and that *hvcmf4* primarily reduces vascular dimensions rather than disrupting the basic rachis vein layout (Rutten et al., 2024). Separately, blocking light exposure to developing spikes, which normally accumulate chlorophyll while still enclosed within leaf sheaths, impairs floral survival and pollen viability, particularly near the inflorescence tip (Babanna et al., 2026). Together, these studies converge on a model in which greening, carbohydrate metabolism, and local vascular support cooperate to sustain developing spikelets. However, how these processes are spatially organized at the resolution of meristematic, vascular-associated, and apical degenerating spike tissues remains unknown.

Here, we used MALDI-MSI to map metabolite distributions across early barley spikelet development in wild-type Bowman and the premature degeneration mutant, *hvcmf4*. By integrating spatial metabolite maps with spike-region transcriptomics and published spatial transcriptome data, we identified metabolite domains associated with reproductive meristem maintenance, vascular support, and the onset of apical degeneration, providing a spatial metabolic framework that helps to understand how local metabolic environments can shape cereal spikelet fate.

## Results

### MALDI-MSI reveals spatial metabolic organization across early inflorescence development in barley

Our previous work established the barley spike PTD as a spatially patterned developmental process and showed that the premature degeneration mutant *hvcmf4* disrupts *HvCMF4*-dependent spike greening, carbohydrate metabolism, vascular development, and hormone balance (Huang et al., 2023; Shanmugaraj et al., 2023b). These findings identified major physiological processes associated with spikelet survival but did not resolve how metabolic states are distributed across meristematic, vascular-associated, and degeneration-prone regions of the developing spike. Because many metabolites are small molecules that can function not only as metabolic intermediates but also as developmental signals, redox-associated compounds, membrane-remodeling factors, and indicators of cellular viability, we used MALDI-MSI as an unbiased spatial survey to build a spatial metabolic framework for early barley inflorescence development.

We analyzed longitudinal sections of wild-type Bowman and *hvcmf4* spikes across stages spanning spikelet initiation, spike growth, and visible PTD, including W3.0, W3.5, W4.5, W5.0, W6.5, and W7.5 (Figure 1A-D). Early stage spikes were analyzed as intact longitudinal sections, whereas later-stage spikes were divided into basal and apical regions because of the increasing spike size and awn complexity. The division was made around node 17, corresponding approximately to the region above which spikelets frequently degenerated in the *hvcmf4* mutants. This design enabled the comparison of basal viable tissues and apical degeneration-prone tissues across genotypes and developmental stages. MALDI-MSI was performed on 10 µm cryosections at 15 µm lateral resolution in positive ionization mode across an *m/z* range of 80-1000 Da, a window selected to preferentially capture a broad spectrum of low-molecular-weight metabolites and lipids. The resulting ion images included features assigned to metabolites, such as amino acids, sugars, and chlorophyll-related compounds, as well as other biologically relevant classes, including polyamines and lipid-associated signals (Supplemental Table 1). Assignments were supported by comparison with authentic standards, with accurate-mass database searches and prior published MSI annotations (Palmer et al., 2017). As expected for untargeted MSI, many detected ion features remained unassigned; however, several of these showed reproducible anatomical patterns, indicating that the spatial distribution itself can provide useful biological information for prioritizing candidate metabolite domains.

**Figure 1.**
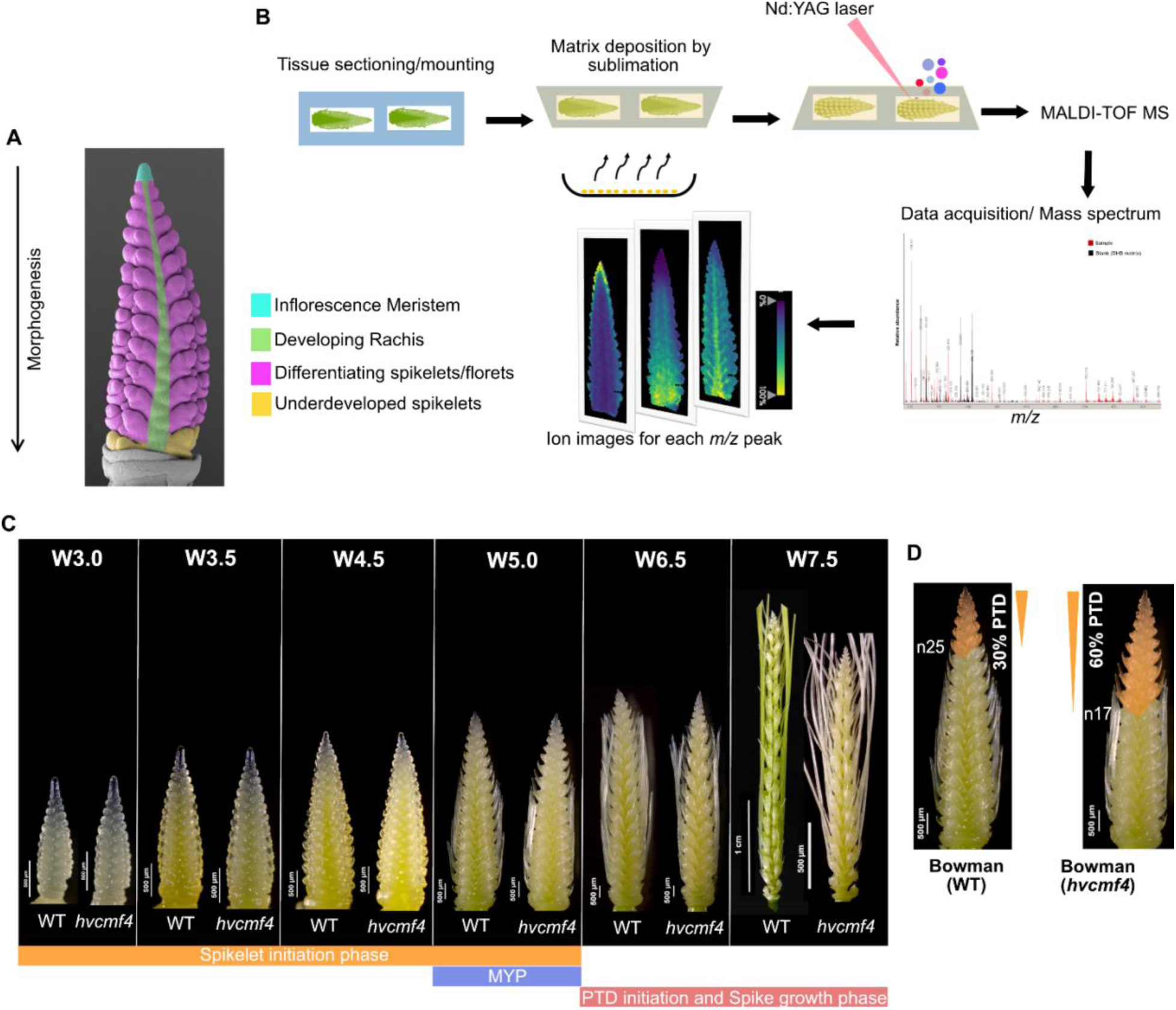
Spatial metabolomics of barley spikelet survival by MALDI-MSI. (A) Schematic of a developing barley spike indicating major anatomical domains analyzed in this study, including the inflorescence meristem, rachis, differentiating spikelets/florets, and underdeveloped apical spikelets. (B) Overview of the MALDI-MSI workflow. Developing spikes were cryosectioned, mounted onto conductive slides, coated with matrix by sublimation, and analyzed by MALDI-TOF mass spectrometry. Ion images were generated for selected m/z features and aligned with optical tissue images. (C) Representative wild-type Bowman and *hvcmf4* spikes across Waddington stages W3.0, W3.5, W4.5, W5.0, W6.5, and W7.5, spanning spikelet initiation, maximum yield potential, PTD initiation, and spike growth phases. (D) Representative W6.5 spikes showing node-based separation of developing and degenerating regions. Wild-type Bowman shows limited apical PTD, whereas *hvcmf4* displays expanded premature degeneration. Scale bars are indicated in the panels. n, rachis node; MYP, Maximum yield potential; W, Waddington scale.

Many ion images showed non-uniform distributions aligned with developmental landmarks, including the inflorescence meristem (IM), spikelet meristems (SMs), spikelet-rachis attachment regions, vascular-associated domains, and apical tissues undergoing degeneration (Figure 2A). Therefore, we first used broadly assigned primary metabolite classes to establish the physiological landscape of the developing spike. Amino acid-, carbohydrate-, and chlorophyll-associated signals were particularly informative because these pathways have previously been linked to developmental PTD, spike greening, and *HvCMF4*-dependent spikelet survival (Huang et al., 2023; Shanmugaraj et al., 2023b). These maps provide a reference framework for distinguishing general metabolic transitions associated with development and degeneration from more spatially restricted signatures associated with meristematic maintenance or vascular support.

**Figure 2.**
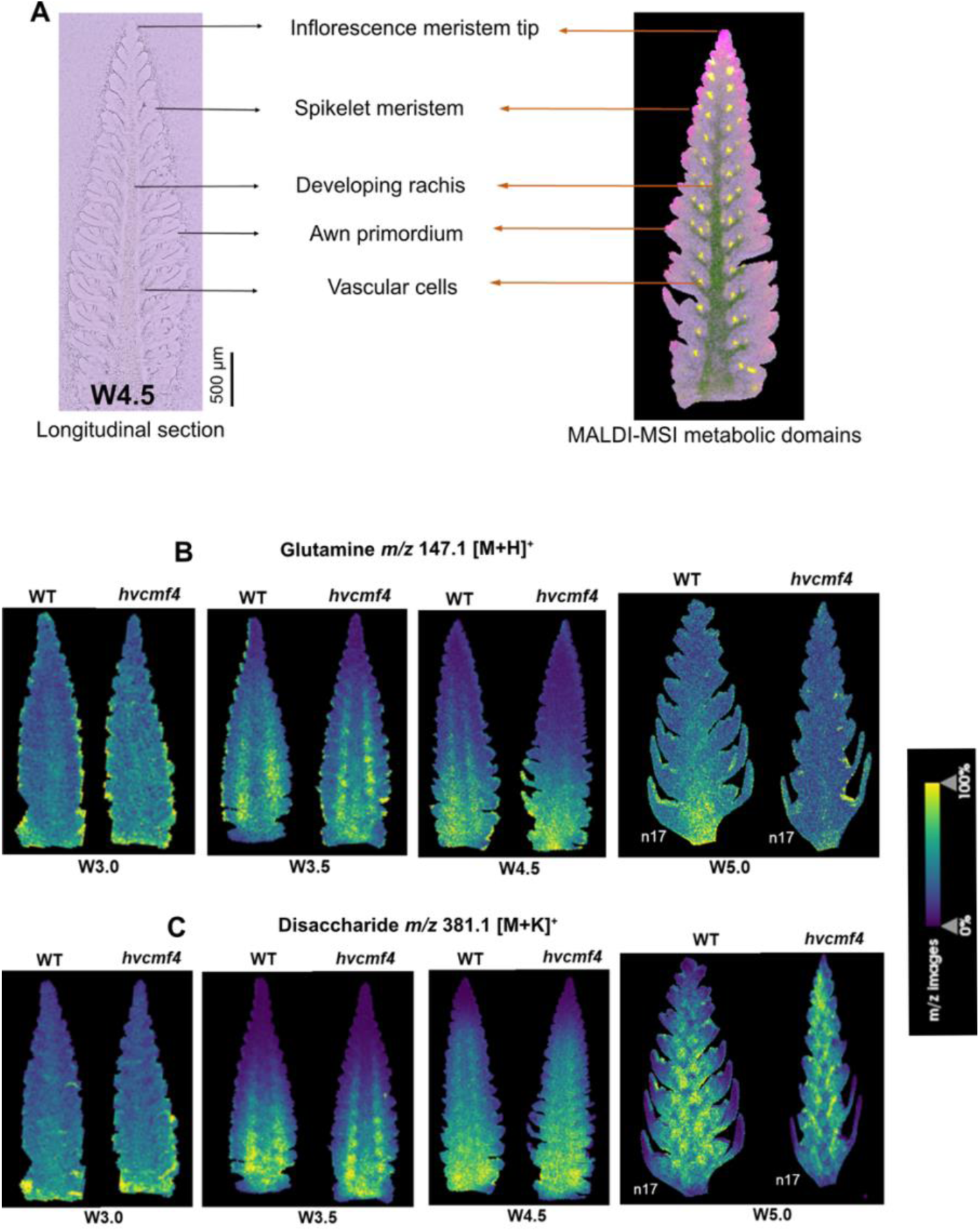
Primary metabolite-associated gradients in developing wild-type Bowman and *hvcmf4* spikes. **(A)** Optical image (left) of a representative W4.5 longitudinal spike section showing major anatomical landmarks used for MSI interpretation, including the inflorescence meristem tip, spikelet meristems, developing rachis, awn primordia, and vascular cells and corresponding multi-ion map from MALDI MSI (right). **(B)** MALDI-MSI ion images showing the distribution of the glutamine-associated signal *m/z* 147.1 [M+H]+ in wild-type Bowman and *hvcmf4* spikes from W3.0 to W5.0. The signal is enriched in basal and central spike regions during early development and becomes reduced in apical regions, with earlier and stronger depletion in *hvcmf4*. **(C)** MALDI-MSI ion images showing the distribution of the disaccharide-associated signal *m/z* 381.1 [M+K]+ in wild-type Bowman and *hvcmf4* spikes from W3.0 to W5.0. The disaccharide-associated signal shows stage-dependent redistribution and stronger accumulation in degeneration-prone mutant apical regions at later stages. Ion images are shown after TIC normalization, and color scales indicate relative normalized ion intensity. Scale bars and node positions are indicated in the panels. n, rachis node.

Amino acid-associated signals revealed a clear spatial transition along the developing spike axis. The glutamine-associated signal *m/z* 147.1 [M+H]^+^ was enriched in the proximal and central spike regions during spikelet initiation and growth, but declined in apical tissues associated with later PTD. In *hvcmf4*, this depletion occurred earlier (W4.5) and extended farther downwards (Figure 2B). Interestingly, this ion remained undetected in the mutant apical sections at W6.5 and W7.5. In contrast, the asparagine-associated signal *m/z* 133.1 [M+H]^+^ accumulated preferentially in apical degeneration-prone tissues and expanded over a broader domain in the mutant, similar to our previous findings during developmental PTD (Figure 2C; Supplemental Figure 1A). These opposing patterns indicate that *hvcmf4* accelerates and spatially expands the nitrogen metabolic transition normally associated with developmental PTD.

Carbohydrate-associated ions also distinguished viable regions from degeneration-prone regions. A disaccharide signal *m/z* 381.1 [M+K]^+^ showed stage-dependent redistribution and increased in apical mutant regions from W5.0 onward, whereas higher-order oligosaccharide-associated signals were detected only in the spike growth phase, preferentially retained in viable spikelet regions, and reduced in degenerating apical nodes (Figure 2D; Supplemental Figure 1B). Because several hexose disaccharides, including sucrose and trehalose, can contribute to the same potassium-adduct mass range, we conservatively refer to this feature as a disaccharide-associated signal rather than assigning it to sucrose alone. This interpretation is also consistent with our previous invasive measurements during developmental PTD, where sucrose was reduced, but trehalose accumulated in apical degeneration-prone tissues (Shanmugaraj et al., 2023b). Thus, the MSI patterns are consistent with the established association between PTD and altered sugar metabolism, but now place these changes within the anatomical context of the developing spike and reveal where carbohydrate-associated signals are retained or depleted in the *hvcmf4*.

We next examined chlorophyll a (*m/z* 893.5 [M+H]^+^)- and pheophytin a (*m/z* 871.5 [M+H]^+^)-associated ions because *HvCMF4* function and pre-anthesis inflorescence greening both implicate local greening in floral survival after organ initiation. In wild-type Bowman, chlorophyll-associated signals followed an acropetal developmental pattern, consistent with the progressive greening of the developing rachis and spikelet-associated tissues. In *hvcmf4*, these signals were reduced in spikelet-associated tissues at later stages, consistent with impaired greening and premature tip degeneration (Supplemental Figure 2).

Together, these high-resolution maps show that primary metabolites and greening are spatially patterned during early barley spike development and are altered in the degeneration-prone apical tissues of *hvcmf4*.

### A spermidine-associated signal defines a reproductive meristem metabolic domain

After establishing broad metabolic gradients across the developing spike, we examined whether more spatially restricted MSI features could define the metabolic states associated with specific developmental domains. Among these features, *m/z* 146.1 showed a strong association with meristematic tissues. At W3.0, this signal accumulated strongly in the inflorescence meristem (IM) of both wild-type Bowman and *hvcmf4* (Figure 3A,B). At W3.5 and W4.5, it remained enriched in the IM and became prominent in SMs, peripheral regions of developing spikelet primordia, and awn primordia. This reproducible pattern suggests that *m/z* 146.1 marks an active reproductive meristem-associated metabolic state during early barley spike development.

**Figure 3.**
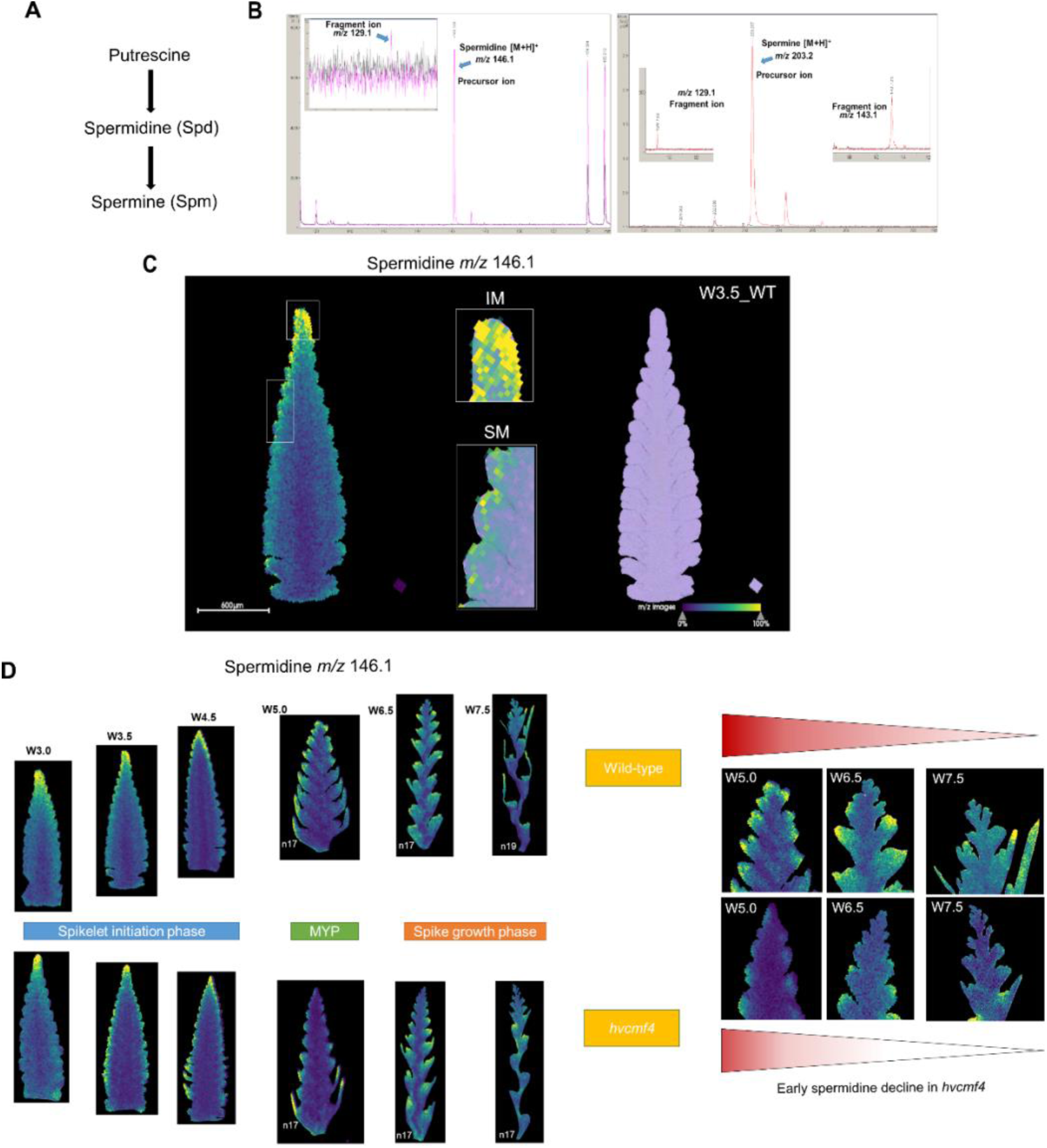
A spermidine-associated signal marks reproductive meristem domains. **(A)** Simplified overview of spermidine and spermine biosynthesis. **(B)** MALDI-TOF MS spectra of a spermidine and spermine standards using DHB matrix. Magenta spectrum – spermidine standard showing a prominent *m/z* 146.1 [M+H]+ signal and a related *m/z* 129.1 fragment. Red spectrum – spermine standard in the right panel highlights the precursor ion of spermine with *m/z* 203.2 [M+H]+ with maximum intensity and two inner mass spectra showing weak fragment ions of spermine at *m/z* 129.1 and *m/z* 143.1. **(C)** MALDI-MSI of wild-type Bowman at W3.5 showing enrichment of the *m/z* 146.1 signal in the inflorescence meristem (IM) and spikelet meristems (SMs). Magnified overlay views with optical image highlight localized accumulation in meristematic domains. **(D)** Developmental series of wild-type Bowman and *hvcmf4* spikes showing the spatial distribution of the m/z 146.1 signal from W3.0 to W7.5. The signal is enriched in meristematic and young spikelet-associated tissues during spikelet initiation and early spike growth. Later-stage apical spike regions showing premature loss of the spermidine-associated signal in degeneration-prone apical tissues of *hvcmf4* compared with wild-type Bowman. Ion images are shown after TIC normalization, and color scales indicate relative normalized ion intensity.

To identify this meristem-associated ion, we compared its mass and fragmentation pattern with authentic standards and barley spike extracts. Database searches, MALDI-TOF MS analysis of a spermidine standard, and analysis of polar spike extracts supported the assignment of *m/z* 146.1 to the spermidine protonated molecular ion [M+H]^+^ (See methods; Supplemental Table 1). MALDI-TOF MS and LC-ESI-QTOF MS analysis using a spermidine standard confirmed its ionization as [M+H]^+^ and in both cases, the formation of the *m/z* 129.1 fragment corresponding to the neutral loss of ammonia [M-NH3+H]+(Figure 3B; Supplemental Figure 3A). In the MSI datasets, low-intensity ions at *m/z* 203.2 and *m/z* 129.1 showed spatial patterns similar to those at *m/z* 146.1 (Figure 3B; Supplemental Figure 3B). Analysis of a spermine standard supported *m/z* 203.2 as a spermine-associated ion, with related fragments overlapping the spermidine-associated mass range (Supplemental Figure 3A). Therefore, we refer to *m/z* 146.1 as a spermidine-associated MSI signal and co-localized *m/z* 203.2 and *m/z* 129.1 features as spermine/polyamine-associated signals.

Because MSI reports metabolite-associated ion distributions but does not directly identify the cells producing these metabolites, we used published barley spatial transcriptome data at W3.5 to test whether the spermidine-associated MSI pattern was supported by the local expression of polyamine biosynthetic genes (Demesa-Arevalo et al., 2026). If *m/z* 146.1 reflects a meristem-associated polyamine state, genes required for decarboxylated S-adenosylmethionine production and spermidine/spermine biosynthesis would be expected to show enriched expression in meristematic domains. Consistent with this expectation, candidate genes, barley *S-ADENOSYLMETHIONINE DECARBOXYLASE (HvSAMDC)*, *SPERMIDINE SYNTHASE (HvSPDS)* and SPERMINE SYNTHASE *(HvSPMS)* were enriched in meristem-associated spatial domains (Supplemental Figure 4; Supplemental Table 2). This independent transcriptomic pattern supports the MSI-based inference that reproductive meristems maintain a local polyamine biosynthesis or maintenance state.

Next, we examined whether this spermidine-associated meristem domain was maintained during spike growth or disrupted in the premature degeneration mutant. At W5.0, wild-type spikes retained a strong *m/z* 146.1 signal in the inflorescence meristem and developing spikelets. In *hvcmf4*, the signal was weaker and less spatially distributed in the apical meristematic regions (Figure 3C). By W6.5 and W7.5, the spermidine-associated signal was strongly reduced in mutant apical nodes prone to PTD, whereas wild-type spikes showed a comparable reduction only later and mainly in the most apical PTD zone. Thus, the premature loss of this meristem-associated spermidine signal in *hvcmf4* preceded or coincided with enhanced apical spikelet degeneration.

Together, MALDI-MSI and published single-cell transcriptome data identified polyamine-associated meristem domains in developing barley spikes. The spatial overlap between spermidine-associated ion accumulation and meristem-enriched expression of polyamine biosynthetic genes suggests that active reproductive meristems maintain a local polyamine-rich environment. The premature depletion of this domain in *hvcmf4* raises the possibility that developmental PTD involves a developmental transition away from meristematic polyamine maintenance, and that this transition is activated earlier in the mutant.

### Premature apical reprogramming of polyamine metabolism in *hvcmf4* resembles an accelerated developmental PTD (dPTD) program

Spermidine is synthesized from putrescine through reactions that depend on decarboxylated S-adenosylmethionine and SPDS activity, whereas spermine is produced from spermidine by SPMS. In contrast, polyamine back-conversion and catabolism are mediated by POLYAMINE OXIDASES (PAOs), which contribute to cellular redox metabolism through hydrogen peroxide production (Blázquez, 2024) (Figure 4A). Given that the spermidine-associated MSI signal marked active reproductive meristems and was depleted in degeneration-prone apices, we examined whether spike PTD was accompanied by a transcriptional shift from polyamine biosynthesis or maintenance toward polyamine oxidation.

**Figure 4.**
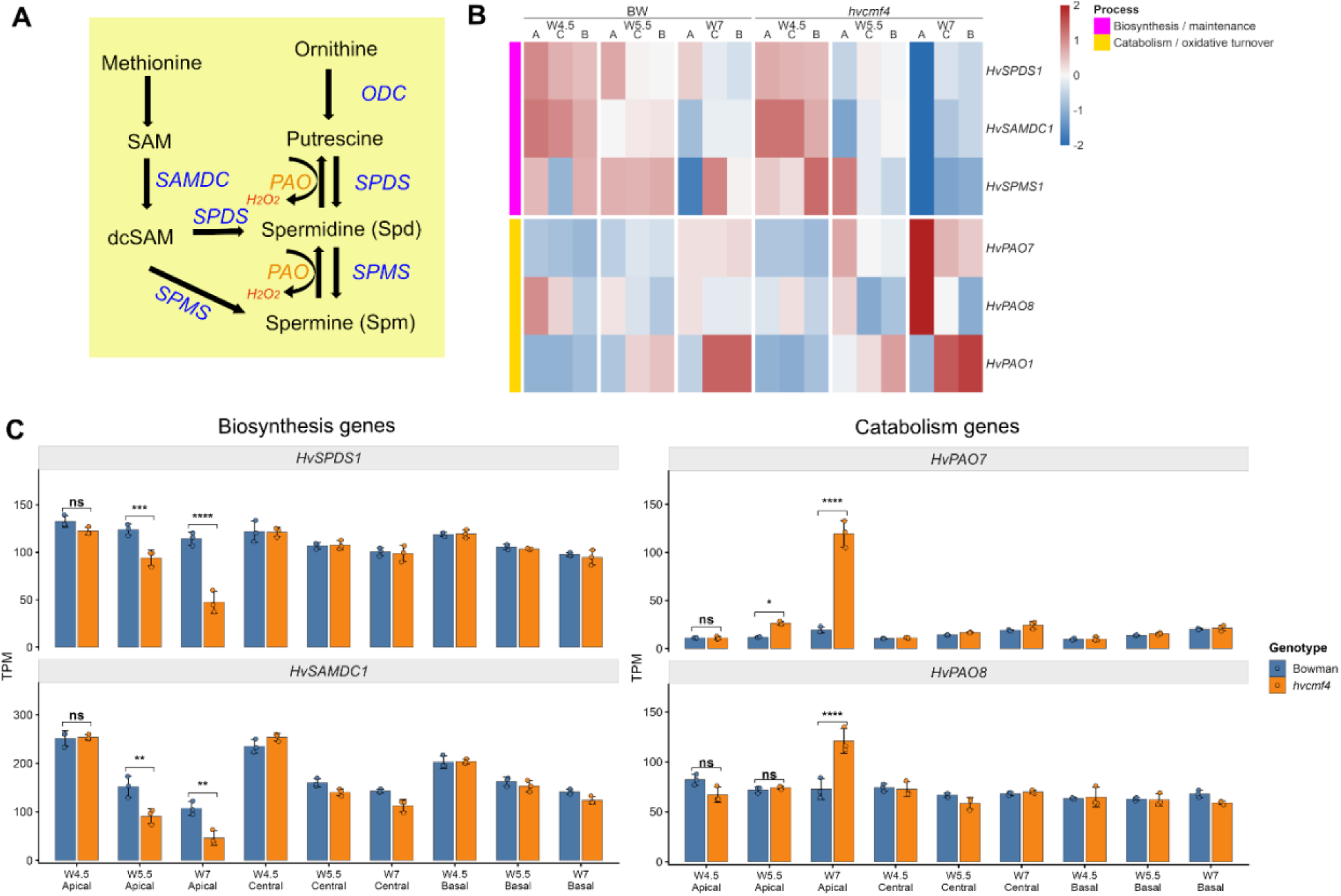
Polyamine pathway expression supports premature activation of a developmental PTD-associated turnover program in *hvcmf4* (A) Simplified pathway schematic showing putrescine, spermidine and spermine biosynthesis and catabolism. (B) Heatmap of selected polyamine-related genes across developmental stages and spike positions in Bowman and *hvcmf4*. Mean TPM values from biological replicates were log2-transformed [log2(TPM + 1)] and row-wise Z-score normalized. (C) Bar plots show TPM values for selected genes, including *HvSPDS1*, *HvSAMDC1*, *HvPAO7* and *HvPAO8*. Error bars indicate SD, and dots represent individual biological replicates. Statistical significance between Bowman and *hvcmf4* in apical samples was determined by two-way ANOVA and Holm-adjusted *P* values, with significance indicated as ns, not significant; P < 0.05, *; P < 0.01, **; P < 0.001, ***; and P < 0.0001, ****. Supplemental Bowman and Morex analyses show that developmental PTD is associated with a late transition from polyamine biosynthesis/maintenance toward PAO/CuAO-mediated oxidative turnover.

To place this transition in the context of regular barley spike PTD, we re-examined Bowman spike transcriptome data across W5.0, W6.5, and W7.5 (Shanmugaraj et al., 2023b). In Bowman, visible spike PTD initiates around W7 or later. We observed that while pre-W7.0 spike tissues retained expression of biosynthesis or maintenance genes, including *HvSAMDC1*, *HvSPDS1* and *HvSPMS1*, in post-W7.0 spikes W7.5 apical or degeneration-associated tissues showed increased expression of selected *PAO/CuAO* genes, including *HvPAO7* and related oxidative-turnover candidates (Supplemental Figure 5; Supplemental Table 4). A comparable developmental logic was observed in the six-rowed cultivar, Morex (Supplemental Figure 5; Supplemental Table 5). Although the timing and individual *PAO/CuAO* genes differed from Bowman, Morex also showed position- and stage-dependent regulation of polyamine biosynthesis and oxidative turnover genes during the PTD window. These data suggest that regular barley spike PTD involves a late transition away from meristem-associated polyamine maintenance toward polyamine oxidation and remodeling, with genotype- or spike architecture-dependent differences in timing and gene usage.

We then asked whether the premature degeneration mutant *hvcmf4* activates this transition earlier in the apical spike tissues than in the wild-type. Using published RNA-seq data from wild-type Bowman and *hvcmf4* spikes, we examined genes associated with spermidine/spermine biosynthesis or maintenance, including *HvSPDS1*, *HvSAMDC1* and *HvSPMS1*, together with polyamine catabolism genes, including *HvPAO7* and *HvPAO8* (Figure 4B; Supplemental Figure 6; Supplemental Table 3). Because PTD is strongest in apical tissues, we compared its expression across the apical, central, and basal spike regions. The clearest genotype-dependent reprogramming occurred in the apex, where *hvcmf4* underwent premature and extended spike PTD. At W5.5, *HvPAO7* was already elevated in the *hvcmf4* apex relative to the wild type, whereas *HvSPDS1* and *HvSAMDC1* were reduced. By W7, this apical shift became more pronounced, with a strong induction of *HvPAO7*, increased expression of *HvPAO8*, and a marked reduction in *HvSPDS1* and *HvSAMDC1* in the mutant apical tissue (Figure 4C). In contrast, genotype-dependent differences in the central and basal regions were weaker or less consistent, indicating that polyamine pathway reprogramming is concentrated in the degeneration-prone apical domain. These expression patterns suggest that *hvcmf4* apices prematurely shift from a spermidine biosynthesis or maintenance state to *PAO*-mediated polyamine oxidation. The early induction of *HvPAO7* at W5.5, before the stronger W7 response, was consistent with the earlier onset and broader extent of spike PTD in the mutant. Thus, viable reproductive meristems are associated with local polyamine maintenance, whereas degeneration-prone *hvcmf4* apices exhibit reduced spermidine biosynthetic gene expression and enhanced *PAO* expression.

Together, the MALDI-MSI, spike-region RNA-seq, and published spatial transcriptome data support a coherent polyamine model for barley spikelet fate. Active IM and SMs maintain a spermidine-associated metabolic state supported by the local expression of polyamine biosynthetic genes. During regular spike PTD in Bowman and Morex, this state is remodeled late as apical tissues transit toward *PAO/CuAO*-mediated oxidative turnover. In *hvcmf4*, the same transition was activated earlier and more strongly in apical spike tissues, coinciding with reduced *HvSPDS1* and *HvSAMDC1* expression and early induction of *HvPAO7*. These findings identify altered polyamine turnover as a molecular feature of degeneration-prone apical spike tissues and suggest *PAO*-mediated oxidation as a candidate process associated with the premature loss of meristematic viability.

### An LPC-associated lipid-remodeling domain marks spikelet supply zones and links to auxin canalization

In addition to the spermidine-associated meristem signal, MALDI-MSI revealed a distinct spatial signature linked to lysophospholipid metabolism. Two co-localized ions, *m/z* 496.3 ([M+H]^+^) and *m/z* 534.3 ([M+K]^+^), showed a reproducible pattern in spikelet attachment and vascular-associated regions (Figure 5A-B). Database searches, expected adduct behavior, targeted LC-MS/MS support and spatial co-localization supported assignment of these features as the molecular protonated ion [M+H]+ and potassium adduct [M+K]+ derived from the ionization of palmitoylated lysophosphatidylcholine, LPC(16:0) (Supplemental Figure 7; See methods).

**Figure 5.**
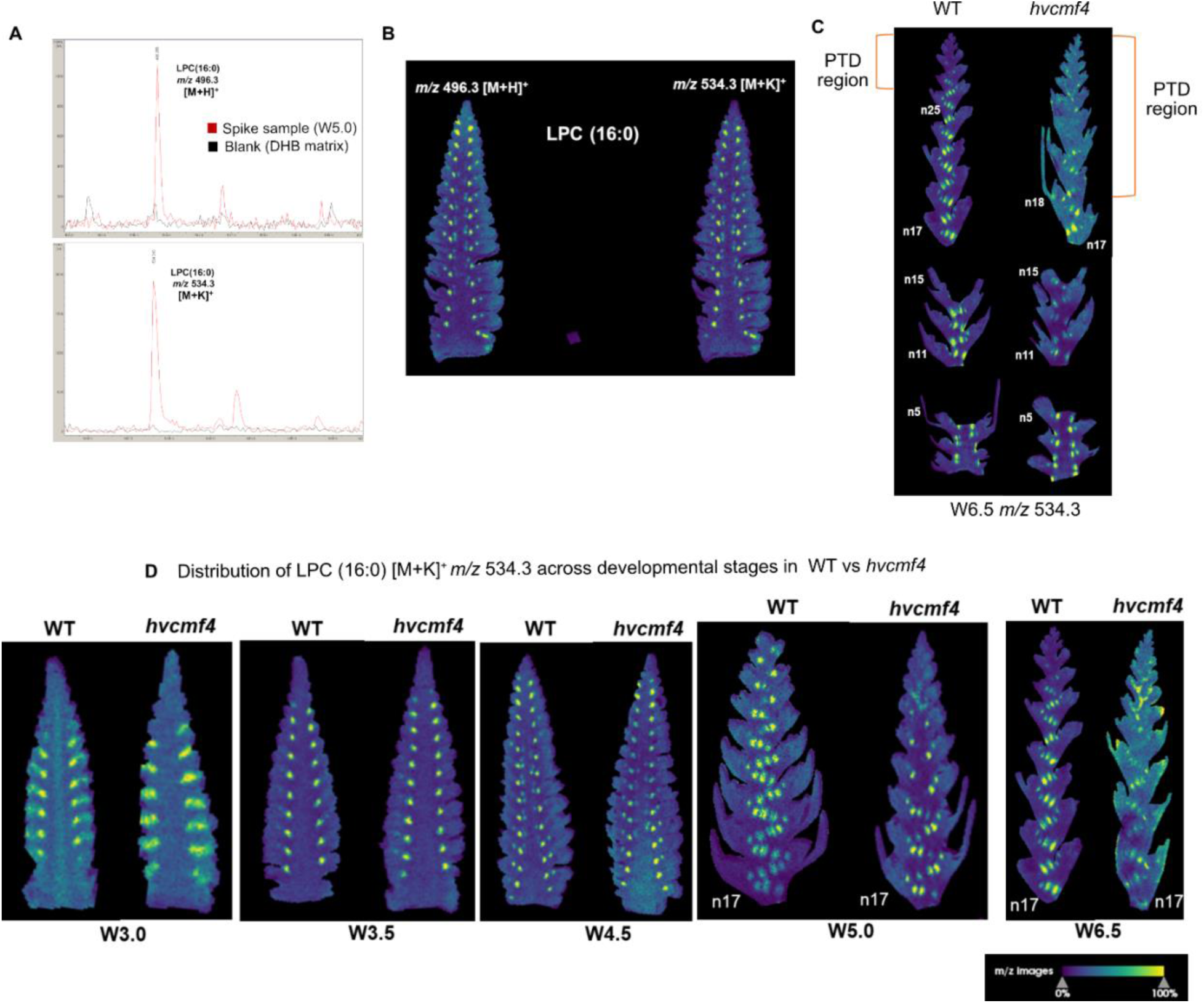
An LPC-associated signal marks spikelet attachment and vascular-associated supply zones. **(A)** Mass spectra from barley spike extracts using DHB matrix showing LPC(16:0)-associated [M+H]^+^ *m/z* 496.3 and *m/z* 534.3 [M+K]^+^ ions. **(B)** Representative MALDI-MSI ion images of wild-type Bowman spikes showing spatial co-localization of *m/z* 496.3 and *m/z* 534.3 signals. The LPC-associated signal accumulates in discrete domains associated with spikelet attachment regions and vascular-like trajectories. **(C)** Comparison of basal and apical spike regions showing reduced or disrupted LPC-associated signal in degeneration-prone apical tissues of *hvcmf4*, whereas clearer vascular-associated signal is retained in viable regions. **(D)** Developmental distribution of the *m/z* 534.3 LPC-associated signal across wild-type Bowman and *hvcmf4* spikes. During early spikelet initiation, the signal forms localized foci and canalized patterns extending from spikelet primordia toward basal attachment regions. During later spike growth, the signal is detected along vascular-associated axes beneath developing spikelets and is retained in viable basal regions. Ion images are shown after TIC normalization, and color scales indicate relative normalized ion intensity.

The LPC-associated signal exhibited a developmental pattern distinct from that of the spermidine-associated meristem domain (Figure 5C). During early spikelet initiation, *m/z* 496.3 showed canalized patterns extending from advanced spikelet primordia to their basal attachment regions. At W3.5, the signal intensified at the boundaries between adjacent spikelet primordia, and by W4.5 additional foci appeared within the developing spikelet primordia and at the interface between the spikelets and the central developing rachis. During later spike growth, *m/z* 496.3 and *m/z* 534.3 signals were detected along two lateral vascular-like axes beneath the developing spikelets, together with weaker foci within the spikelet tissues (Figure 5C). As lysophosphatidylcholine (LPC) is a bioactive lysophospholipid implicated in plant signaling, membrane activity, and phospholipid homeostasis, we reasoned that this spatial feature could mark a regulated lipid-remodeling domain associated with spikelet support rather than a passive lipid distribution (Drissner et al., 2007; Poulsen et al., 2015).

While this LPC-associated domain becomes progressively weaker in degeneration-prone tissues, this decline was more pronounced in the *hvcmf4* mutant. At W5.0 and W6.5, *m/z* 496.3 and *m/z* 534.3 signals were strongly reduced or absent in apical mutant spike nodes, whereas basal viable regions retained clearer vascular-associated signals. In wild-type Bowman, comparable loss of the LPC-associated pattern was largely restricted to later-degenerating apical spike nodes (Figure 5C; Figure 6A). Multi-ion overlays showed that the LPC-associated signal was spatially distinct from the chlorophyll-associated signals, indicating that lysophospholipid-associated patterning and greening-associated metabolism represent separable components of the spike metabolic landscape (Figure 6B). This distinction is important because previous anatomical studies have shown that *hvcmf4* reduces rachis vascular dimensions without abolishing the basic vascular layout, suggesting that local vascular support may be compromised at the level of tissue maturation, lipid remodeling, or transport function (Rutten et al., 2024).

**Figure 6.**
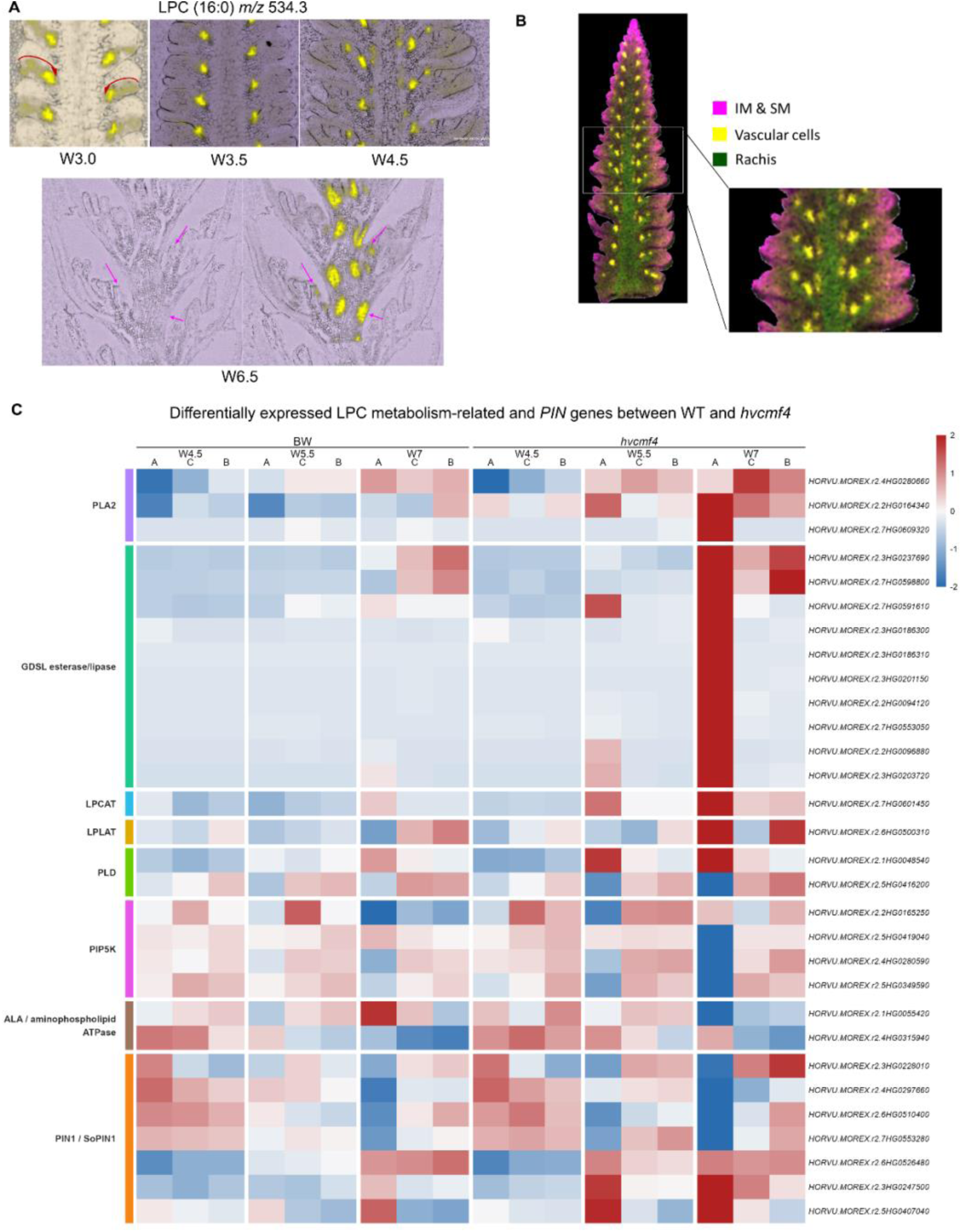
Lipid-remodeling and auxin transport-associated genes support the LPC-associated vascular domain. **(A)** Optical images and MALDI-MSI ion images showing the spatial relationship between the LPC-associated signal and spikelet attachment or vascular-associated regions. Yellow signal indicates the LPC(16:0)-associated ion m/z 496.3, highlighting localized accumulation near spikelet bases and vascular-like trajectories. **(B)** Multi-ion overlay of representative barley spike sections showing that the LPC-associated signal occupies spatial domains distinct from chlorophyll- and pheophytin-associated greening signals. The overlay supports separation of lysophospholipid-associated patterning from general greening-associated metabolism. **(C)** Expression heatmap of candidate genes associated with phosphatidylcholine turnover, LPC production and recycling, membrane lipid signaling, lipid asymmetry, and auxin transport. Gene groups include PLA2-like, GDSL ESTERASE/LIPASE, LPCAT/LPLAT, PLD, PIP5K, ALA/P4-ATPase, and PIN1/SoPIN-associated candidates. Expression values are shown across developmental stages, spike regions, and genotypes. The strong stage-, position-, and genotype-dependent regulation of these genes supports the presence of a regulated lipid-remodeling program associated with spikelet attachment and vascular-associated supply zones. Color scale indicates row-scaled expression values.

The biological significance of this LPC-associated pattern is supported by previous studies showing that LPC can act as a bioactive lysophospholipid signal in plants. In arbuscular mycorrhizal symbiosis, LPC species, including LPC(16:0), induce rapid cellular responses and mycorrhiza-responsive phosphate transporter expression (Drissner et al., 2007; Vijayakumar et al., 2016). Although the developmental context is different, these findings show that LPC species can function as plant signaling lipids at biologically active interface domains. We therefore asked whether loss of the organized LPC-associated MSI domain in *hvcmf4* was accompanied by transcriptional reprogramming of lipid-remodeling pathways. We examined the expressed candidate genes associated with phosphatidylcholine turnover, LPC production, lysophospholipid recycling, membrane lipid signaling and lipid asymmetry. These included *PHOSPHOLIPASE A2 (PLA2)*-like genes, *GDSL ESTERASE/LIPASE* genes, *LYSOPHOSPHATIDYLCHOLINE (LPCAT)/ LYSOPHOSPHOLIPID (LPLAT) ACYLTRANSFERASES*, *PHOSPHOLIPASE D (PLD)*, *PHOSPHATIDYLIONOSITOL 4-PHOSPHATE 5-KINASE (PIP5K),* and *AMINOPHOSPHOLIPID (ALA)/P4-ATPase* candidates. These genes showed strong stage-, position- and genotype-dependent expression, indicating that the developing barley spike contains a regulated lipid-remodeling program (Figure 6C; Supplemental Table 6). In *hvcmf4*, many lipid-remodeling genes, particularly *GDSL ESTERASE/LIPASE* candidates, were strongly altered in apical spike tissues during W5.5-W7, coinciding with loss of the organized LPC-associated MSI pattern. Thus, premature apical floral and spike degeneration is not associated with the simple absence of lipid metabolism, but with the misregulation of a spatially organized lipid-remodeling state.

The dynamics and geometry of the LPC signal further suggest a link to auxin-guided vascular patterning. Auxin transport contributes to primordium initiation, vascular trace formation, and the connection of new organs to existing vascular tissues. In grasses, the *SISTER OF PIN-FORMED1 (SoPIN1)*-like auxin efflux carrier gene is associated with auxin maxima and organ positioning, whereas internal *PIN1a/PIN1b*-like activity is linked to auxin canalization and vein patterning (O’Connor et al., 2014; O’Connor et al., 2017). The LPC signals observed in the current study, extending from spikelet primordia toward the basal attachment regions and later along the vascular-like axes, resemble the expected geometry of a local auxin-guided supply domain. Consistent with this interpretation, several *PIN1/SoPIN* candidates and membrane polarity-associated lipid genes showed positional and genotype-dependent regulation (Figure 6C; Supplemental Table 6). Together with the previously observed apical IAA accumulation and reduced vascular development in *hvcmf4*, these patterns suggest that auxin is present in mutant spike apices but may not be effectively converted into stable canalized transport or vascular support.

Next, we used published barley single-cell transcriptome data to evaluate whether lipid-remodeling and auxin transport candidates occupy domains consistent with the LPC-associated MSI pattern (Demesa-Arevalo et al., 2026). Candidate lipid-remodeling genes *PLA*, *LPCAT*, *PLD,* and *PATATIN-like*, together with *PIN* genes, were expressed in regions related to the spikelet meristem base or spikelet attachment-like domains (Supplemental Figure 8). The overlap or close adjacency of lipid remodeling and auxin transport domains provides independent spatial support for the hypothesis that the LPC-associated MSI signal marks a vascular-associated lipid-remodeling domain linked to auxin-guided supply-zone formation. As *PIN*-mediated auxin transport depends on membrane-embedded exporters whose localization and activity are influenced by membrane trafficking and lipid environment, the coordinated reprogramming of lipid-remodeling and *PIN-*associated genes provides a plausible connection between LPC-associated lipid remodeling and auxin canalization (Lee et al., 2010).

Together, these results identify an LPC-associated domain that marks spikelet attachment and vascular-associated supply zones in the developing spike. This domain progressively weakens in degeneration-prone apical spike tissues during development and is prematurely disrupted in *hvcmf4*. In the context of *HvCMF4*-dependent greening, rachis vascular anatomy, lipid-remodeling gene expression and auxin transport-associated gene regulation, these findings support a model in which lysophosphatidylcholine remodeling may help establish or report the local vascular support environment required for proper spikelet development.

## Discussion

### Spatial metabolomics reveals domain-level organization of barley spikelet fate

Spikelet survival in cereals depends on the ability of young reproductive primordia to maintain meristematic activity, establish local vascular support, and avoid premature degeneration. Previous genetic, anatomical, and multi-omic studies of the barley *hvcmf4* mutant established *HvCMF4* as a regulator of spikelet survival through effects on rachis greening, plastidial metabolism, carbohydrate utilization, hormone homeostasis, and vascular development (Huang et al., 2023). Studies on pre-anthesis inflorescence greening further showed that chlorophyll accumulation in sheathed barley spikes supports floral survival after organ initiation, particularly in apical spike regions (Babanna et al., 2026). These findings, together with *HvCMF4/hvcmf4* and rachis vascular studies, point to greening, energy metabolism, and vascular development as interconnected support systems for barley floral fate. However, the spatial organization of these support systems across meristematic, vascular-associated, and degeneration-prone spike tissues remains unclear.

In the current study, MALDI-MSI revealed that barley spikelet fate is associated with anatomically distinct metabolic domains. Primary metabolite and chlorophyll-associated signals formed broad developmental gradients along the spike axis, whereas spermidine- and LPC-associated signals occupied more restricted meristematic and vascular-associated domains, respectively (Figure 2,3,5). The premature loss or disruption of these domains in *hvcmf4* suggests that the enhanced spike PTD in barley is not simply a consequence of global metabolic decline, but reflects the accelerated collapse of local metabolic states that normally support reproductive meristem maintenance and proper spikelet development (Figure 5,6). Integration with RNA-seq and published scRNA-seq transcriptome data strengthened these assignments by linking the spermidine-associated signal to meristem-enriched polyamine biosynthesis and the LPC-associated signal to lipid-remodeling and auxin transport-associated domains (Huang et al., 2023; Demesa-Arevalo et al., 2026) (Figure 6; Supplemental Figure 4,8). Thus, spatial metabolomics converts previously described bulk metabolic defects into domain-level hypotheses regarding developmental function.

### Polyamine turnover as a meristem viability module

The spermidine-associated MSI signal identified a polyamine-rich state in active IM and SM (Figure 3). This is biologically plausible because polyamines, such as putrescine, spermidine, and spermine, are associated with cell proliferation, stress protection, floral development, delayed senescence, and redox metabolism (Blázquez, 2024). Their cationic properties allow them to interact with nucleic acids, membranes, and proteins, whereas their oxidation can generate hydrogen peroxide and connect polyamine metabolism to redox signaling (Murray Stewart et al., 2018). A local spermidine-associated state in reproductive meristems may therefore contribute to growth competence, redox buffering, or cellular viability during spikelet initiation and early spike growth. More broadly, this domain may represent a meristem viability module in which polyamine biosynthesis, macromolecular stabilization, and controlled redox metabolism together create a growth-permissive environment for reproductive meristem cells. In this view, the loss of the spermidine-associated signal is not simply a decline in one metabolite but marks a spatial transition away from a meristem-maintaining metabolic state toward oxidative turnover and developmental degeneration.

The comparison between the regular wild-type developmental spike PTD and *hvcmf4*’s premature degeneration is important for interpreting this module. In wild-type Bowman and the six-rowed cultivar Morex, the apical spike PTD is associated with a late transition from polyamine biosynthesis or maintenance toward *PAO/CuAO*-mediated oxidative turnover (Supplemental Figure 5,6). This suggests that polyamine oxidation is part of a concerted developmental transition during apical remodeling, rather than a process necessarily associated with tissue failure. During dPTD, controlled *PAO/CuAO* activity may participate in apical remodeling, redox signaling, nutrient remobilization, or orderly termination of meristematic growth. In *hvcmf4*, however, this transition was activated earlier and more strongly in the apical tissues, coinciding with the loss of the spermidine-associated meristem signal and reduced expression of spermidine biosynthetic genes (Figure 4). This interpretation is consistent with recent studies linking polyamine maintenance to reproductive fertility in grasses under stress. In wheat, exogenous spermidine reduces drought-induced floret degeneration by mitigating oxidative damage and maintaining energy homeostasis in spikes (Li et al., 2024). *PAO*-mediated oxidation can generate hydrogen peroxide, and redox imbalance has been implicated in apical spikelet degeneration in wheat mutants and stress-induced floret abortion (Du et al., 2026). In rice, spermidine application before heading improves spikelet fertility under heat stress, enhances antioxidant capacity, and is associated with higher endogenous spermidine and spermine levels (Karwa et al., 2022). These studies support a broader role for spermidine and spermine in protecting the developing grass reproductive tissues. Our study extends this concept by demonstrating that a spermidine-associated metabolic state is spatially enriched in barley reproductive meristems and is prematurely depleted in degeneration-prone apical tissues.

The early induction of *HvPAO7* in *hvcmf4* provides a potential link between polyamine turnover and the redox/hormonal imbalance associated with spikelet degeneration. However, this should not be interpreted as evidence that *PAO* activity causes PTD. *HvCMF4* affects multiple interconnected processes, including greening, carbohydrate metabolism, vascular development, and hormone balance (Huang et al., 2023). Instead, premature *PAO* induction may represent a route by which altered polyamine turnover contributes to a broader degeneration-prone state. Notably, *HvPAO7* and *HvPAO8* appear to be tandem paralogs in the barley genome, raising the possibility that a duplicated *PAO* module contributes to the stage-specific regulation of polyamine oxidation during apical spikelet degeneration. In this model, *HvPAO7* may represent an early apical degeneration-associated *PAO*, whereas *HvPAO8* may reinforce polyamine catabolism at later stages of degeneration. Functional analysis is required to determine whether these *PAOs* actively promote degeneration or mark a broader oxidative-remodeling state associated with PTD.

### LPC-associated lipid remodeling links vascular support and auxin canalization

The LPC-associated MSI signal defines a spatially distinct domain from the spermidine-rich meristem. Its localization to spikelet attachment regions and vascular-like trajectories places it in the anatomical context of local spikelet support (Figure 5,6). This is highly relevant to the *HvCMF4* framework because *hvcmf4* reduces vascular development toward the spike apex, while defects in greening and carbohydrate metabolism likely limit the supply available to developing spikelets. Anatomical analyses of barley rachis vasculature suggest that spike fertility is more strongly associated with vascular area and vein size than with vein number, implying that vascular capacity and local supply may be more important than the presence or absence of the basic vascular layout (Rutten et al., 2024). The LPC-associated domain appears in precisely the regions expected to support spikelet attachment and local supply, and its premature disruption in *hvcmf4* provides spatial metabolite evidence of compromised vascular-associated support. This interpretation is further supported by a recent study on barley that identified *HvGELP96*, which encodes a GDSL domain-containing protein, as a regulator of spikelet fertility and grain number. Because *GDSL ESTERASE/LIPASE* candidates were among the lipid-remodeling genes that were most strongly altered in degeneration-prone *hvcmf4* apical spike tissues, these findings provide barley-specific support for a functional connection between lipid-remodeling genes, spikelet development and fertility (Shen et al., 2023).

LPC should not be viewed as a passive membrane breakdown product. LPC has been characterized as a bioactive lysophospholipid signal in plant systems, including arbuscular mycorrhizal symbiosis, in which LPC species induce rapid cellular responses and mycorrhiza-responsive gene expression (Drissner et al., 2007; Poulsen et al., 2015; Wojciechowska et al., 2024). LPC can also influence plasma membrane activity, as shown by the stimulation of ATP-dependent proton accumulation in isolated oat root plasma membrane vesicles (Palmgren and Sommarin, 1989). More recent studies have further suggested that LPC homeostasis through the Lands cycle is important for Arabidopsis root growth and that LPC-binding regulatory proteins can connect lysophospholipid status to the transcriptional control of phospholipid metabolism (Wang et al., 2026). Although barley spike development is a different biological context, these studies support the idea that LPC-associated signals can participate in membrane activity, lipid homeostasis and regulatory responses. In barley spikes, the LPC-associated signal appears at an interface-like region: the spikelet attachment and local vascular supply zone. This raises the possibility that the LPC-associated domain may contribute to, or report, localized membrane remodeling and lipid signaling required for spikelet support (Scherer and André, 1989; Yi et al., 1996; Dong et al., 2014).

The connection between LPC-associated lipid remodeling and auxin canalization provides a mechanistic framework for understanding this domain. Auxin transport is central to organ initiation, vascular trace formation, and the connection of new primordia to existing vascular tissues. In grasses, SoPIN1-like activity is associated with auxin maxima and organ positioning, whereas internal PIN1a/PIN1b-like activity is associated with auxin canalization and vein patterning (O’Connor et al., 2014). Lipid-remodeling enzymes have also been associated with auxin responses. Phospholipase A activity and protein kinase signaling have been implicated in auxin-induced corn coleoptile elongation, and patatin-related phospholipases can influence auxin-responsive cell morphology and organ size (Yi et al., 1996; Dong et al., 2014). Together with evidence that phospholipase activity can affect PIN plasma membrane localization, these studies support a broader link between lipid remodeling, auxin response, and polarized growth. The LPC-associated signal in barley spikes, extending from the spikelet primordia toward the basal attachment regions and later along the vascular-like axes, may therefore mark a lipid-remodeling environment associated with auxin-guided vascular supply zone formation. Imputed scRNAseq transcriptome data further support this interpretation by placing lipid-remodeling and auxin transport-associated genes in the meristem base, primordium-associated or spikelet attachment-like domains (Demesa-Arevalo et al., 2026).

Previous hormonal measurements in *hvcmf4* apices did not appear to be simply auxin-deficient; rather, IAA accumulated in apical spike tissues as vascular defects and premature degeneration emerged (Huang et al., 2023). This suggests that spikelet failure may reflect defective auxin canalization or auxin-response implementation, rather than low auxin abundance alone.

Phospholipid-mediated membrane remodeling provides a plausible cellular link because PIN proteins are membrane-embedded auxin exporters whose localization and activity depend on membrane trafficking and lipid environment (Lee et al., 2010). Evidence from *Arabidopsis* roots indicates that phospholipase activity can influence PIN plasma membrane localization, although this does not demonstrate that LPC directly regulates PIN trafficking in barley spike. Therefore, we propose that the LPC-associated domain may support a lipid-remodeling environment associated with PIN-mediated auxin canalization, vascular maturation, and spikelet attachment zone development. This model helps explain why elevated IAA levels and reduced vascular development can coexist in *hvcmf4*: auxin may be present but not effectively converted into stable canalized vascular support.

### A multilayered metabolic model for barley spikelet survival

Together, our findings support a multilayered model in which barley apical spikelet survival depends on the coordinated maintenance of spike greening, primary metabolism, a polyamine-rich meristematic state, and a lysophospholipid-associated vascular support domain (Figure 7). In wild-type spikes, *HvCMF4*-dependent greening and carbohydrate metabolism provide a supportive layer for developing spikelets. Superimposed on this layer, a spermidine-associated domain marks active reproductive meristems, whereas an LPC-associated domain marks spikelet attachment and vascular-associated supply zones. During regular apical degeneration, these domains are progressively remodeled. In *hvcmf4*, the same transitions are accelerated and spatially expanded in apical tissues, leading to the premature loss of the spermidine-associated meristem state, activation of *PAO*-mediated polyamine turnover, and disruption of the LPC-associated vascular lipid domain.

**Figure 7.**
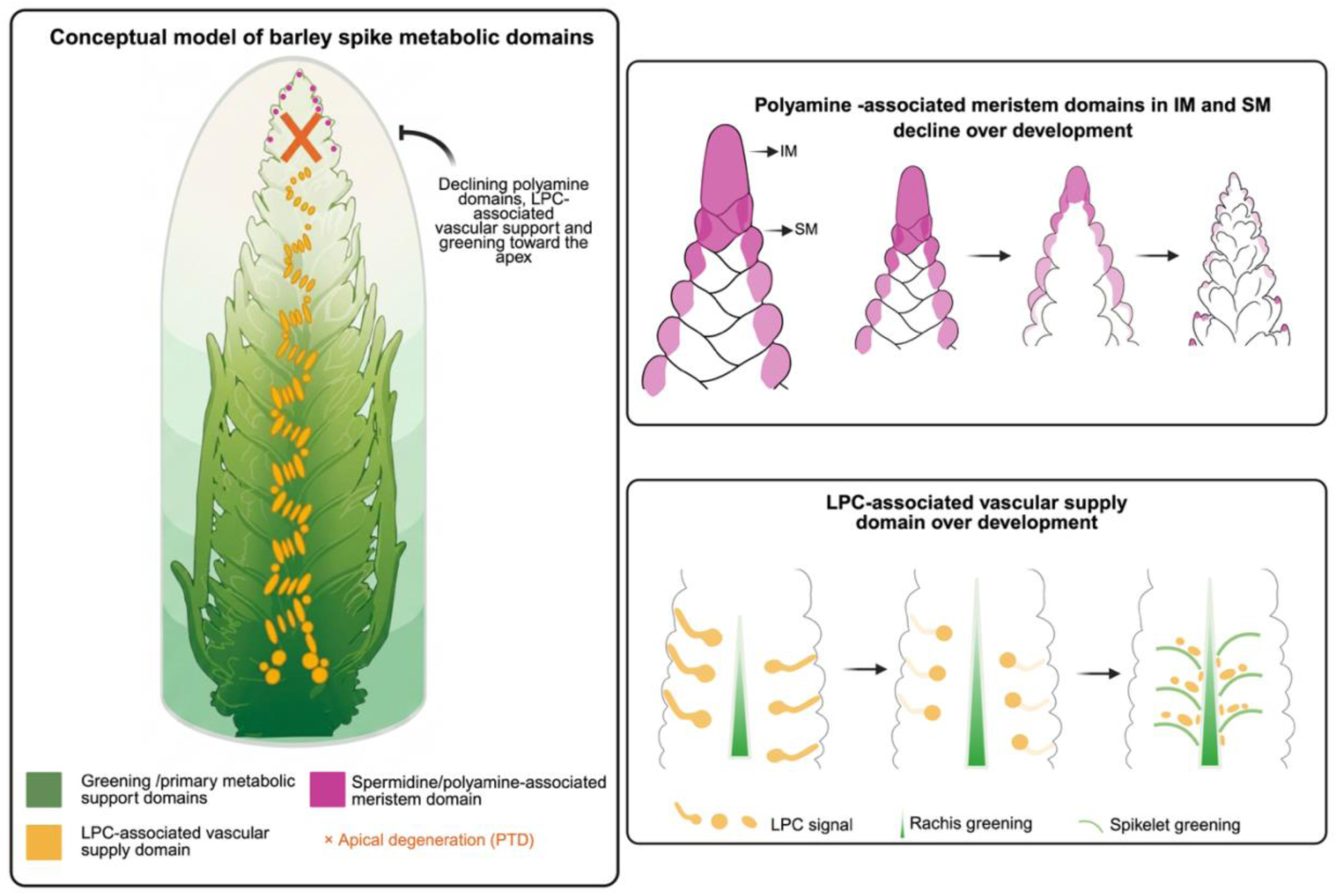
Conceptual model of spatial metabolic domains associated with barley spikelet survival and degeneration. The model summarizes spatial metabolic domains identified during barley spike development. Greening and primary metabolic support are strongest in basal and central spike regions and decline toward the apical tip. Superimposed on this broad metabolic gradient, spermidine/polyamine-associated signals mark reproductive meristem domains, including the inflorescence meristem (IM) and spikelet meristems (SMs), and progressively decline over development. In parallel, LPC-associated signals initially appear as localized domains near spikelet attachment sites and become organized into canalized vascular-associated trajectories that connect developing spikelets with the rachis. Toward degeneration-prone apical tissues, this LPC-associated support domain becomes reduced or fragmented. Together, the model proposes that barley spikelet survival depends on coordinated maintenance of greening/primary metabolism, polyamine-rich meristematic domains, and canalized LPC-associated vascular support domains (created using Biorender.com).

A central contribution of this study is that these relationships become visible only in a spatial context. Bulk metabolite or transcript profiles can identify genotype-level defects in carbohydrate metabolism, greening, lipid remodeling, or polyamine turnover, but they cannot determine whether these changes occur in active meristems, vascular supply zones, or degenerating apical tissues. MALDI-MSI resolved these differences anatomically, whereas transcriptome integration linked metabolite domains to plausible biosynthetic and regulatory programs. This cross-modal framework does not yet establish causality, but it converts metabolite patterns into testable developmental hypotheses.

Several questions have become experimentally tractable. Manipulating polyamine turnover, especially *HvPAO7* activity, could be used to test whether premature PAO activation contributes to the timing or extent of apical spikelet degeneration. Conversely, maintaining spermidine biosynthetic capacity through *HvSPDS1* or *HvSAMDC1* could be used to test whether the meristem-associated polyamine state prolongs apical spikelet viability. For the LPC-associated domain, functional analysis of *PLA2-LIKE*, *GDSL*, *LPCAT/LPLAT,* or *PATATIN-LIKE* genes could determine whether lysophospholipid-associated remodeling actively supports vascular supply or tissue remodeling during degeneration.

Beyond barley, this study illustrates how spatial metabolomics can reveal hidden metabolic domains that are difficult to infer from bulk profiling methods. The regulatory potential of many metabolites depends on their space and time. By integrating MALDI-MSI with mutant analysis, spike-region transcriptomics, and published spatial transcriptome resources, this study provides a framework for discovering metabolic states that influence meristem maintenance, vascular support, and developmental fate in plant reproductive tissues.

## Materials and methods

### Plant material and spike staging

Wild-type barley (*Hordeum vulgare* cv. Bowman) and the near-isogenic *hvcmf4* mutant were grown in 9-cm square pots in a plant chamber (IPK, Gatersleben) with day/night temperatures of 12°C/8°C and 12-h/12-h light–dark photoperiod with a light intensity of 300*µ*E at 80% relative humidity as described previously. Developing spikes were staged using the Waddington scale. Spikes were collected at W3.0, W3.5, W4.5, W5.0, W6.5 and W7.5, covering early spikelet initiation, spikelet primordium development, spike growth and the onset of visible pre-anthesis tip degeneration (PTD). Independent biological samples were collected for MALDI-MSI for each developmental stage and genotype. Additional stage-matched samples were collected for metabolite validation.

### Tissue embedding and cryosectioning

For the early developmental stages W3.0, W3.5, and W4.5, whole-spike longitudinal sections were analyzed. For W5.0, W6.5, and W7.5, spikes were divided into basal and apical regions because of the increasing spike size and awn complexity. The division was made at approximately node 17, corresponding to the region above which spikelets frequently degenerate in the *hvcmf4*. Wild-type and mutant spikes were processed using the same orientation, sectioning, and region division strategy to enable stage-matched comparisons. Tissue embedding and sectioning were performed as previously described (Shanmugaraj et al., 2023a). Briefly, developing spikes were excised, immediately frozen in liquid nitrogen, and stored at −80°C until sectioning. The samples were embedded in freshly prepared 10% (w/v) gelatin. Longitudinal sections of 10 µm thickness were prepared using a CryoStar NX70 cryostat (Thermo Scientific, Germany), with the specimen holder and blade maintained at approximately −10°C and −9°C, respectively. The sections were thaw-mounted onto indium tin oxide-coated glass slides and placed in a vacuum desiccator for approximately 30 min before optical imaging and matrix application.

#### Matrix application

The 2,5-dihydroxybenzoic acid (DHB) matrix was sublimated over the mounted tissue sections using a lab-made sublimator, as described previously (Shanmugaraj et al., 2023a).

#### MALDI-MSI acquisition

MALDI-MSI was performed using an UltrafleXtreme MALDI-TOF/TOF mass spectrometer equipped with a Smartbeam II laser (Bruker Daltonics, Bremen, Germany) and operated in the positive reflectron mode. Data were acquired over an *m/z* range of 80-1000 Da and a sample rate of 1 Gs s^−1^ with a lateral raster step of 15 µm. This mass range was selected to capture low-molecular-weight primary metabolites and additional lipid-associated features. Each raster position was measured in reflectron mode using 600 laser shots at a repetition rate of 1000 Hz. The laser intensity was adjusted to obtain stable analyte signals while minimizing tissue damage beyond each raster position. Images were acquired in the random walk mode. The instrument was calibrated using a polyethylene glycol mixture (PEG) (1:1 PEG 200 and 600, diluted 1:300 in 30% (*v/v*) acetonitrile and 0.1% (*w/v*) trifluoroacetic acid) before each experiment. Optical images were acquired using a digital microscope (Keyence VHX-5000) before MSI and used for anatomical comparisons.

#### MSI data processing and visualization

Reduced MALDI-MSI spectra in the .*mis* format was imported into SCiLS Lab 2021c pro software (Bruker Daltonics) and processed according to the manufacturer’s instructions. Automatic mass-axis alignment was performed using the default constant and equidistant settings. The spectral baselines were corrected using a convolution algorithm with a peak-width parameter of 20. Ion intensities were normalized to the total ion count (TIC) of each spectrum, and ion images were generated using a mass tolerance of [±0.1 Da] around each selected *m/z* value. Optical images acquired before matrix application were co-registered with the MSI datasets and used to identify developmental landmarks and to define regions of interest. Identical processing, normalization, and visualization parameters were applied within stage-matched genotype comparisons.

#### Metabolite annotation and validation

Candidate metabolite assignments were based on observed *m/z* values, expected adduct behavior, comparison with public metabolite databases, METASPACE searches where applicable, previously published plant MSI datasets, and analysis of authentic standards where available (Palmer et al., 2017). Commercial standards of selected amino acids, disaccharides (sucrose and trehalose), spermidine and spermine were prepared at 1 mM and spotted onto AnchorChip targets with DHB matrix. Polar extracts were prepared from manually dissected spike samples from 1 mg of ground frozen tissue powder using 100 µl of 50% (v/v) methanol and analyzed by MALDI-TOF MS to compare tissue-derived ions with standard-derived signals. Selected metabolite and lipid features were further evaluated via RP-UPLC ESI-QTOF MS (Reversed Phase Ultra Performance LC-Electrospray Ionization-Ultra-High-Resolution-Quadrupole Time Of Flight MS) using an Acquity UPLC system (Waters, Germany) coupled to a maXis Impact ESI-QTOF MS (Bruker Daltonik GmbH, Germany) as previously described (Garibay-Hernández et al., 2021). Amino acids and polyamines were confirmed using a commercial standards by retention time, exact mass (error < 3 ppm), isotopic pattern, and MS/MS fragmentation. Metabolite names were reported according to their annotation confidence. Features supported by authentic standards and matching mass and/or fragmentation patterns were reported as assigned metabolites.

#### Transcriptome integration

Published RNA-seq data from wild-type Bowman and *hvcmf4* developing spikes were used to examine the expression of genes involved in polyamine biosynthesis, polyamine catabolism, lipid remodeling, and auxin transport (Huang et al., 2023). Previously reported barley polyamine pathways genes were used as a reference to identify the corresponding gene models in the MorexV2 reference (Monat et al., 2019; Tanwar et al., 2022). Furthermore, candidate barley genes were identified based on functional annotation, and barley reference genome annotation and corresponding gene IDs are listed in Supplemental Table 2. Differential expression values for the analyzed genes are listed in Supplemental Table 3. Expression heatmaps were generated from TPM values after log2(TPM + 1) transformation and row-wise z-score scaling. Bar plots show the mean TPM values with standard deviation and individual biological replicates. Statistical significance between Bowman and *hvcmf4* within each apical developmental stage was assessed after two-way ANOVA and displayed using Holm-adjusted *P* values. Previously published Bowman and Morex PTD transcriptome datasets were re-examined to place polyamine pathway expression in the context of developmental PTD (Supplemental Table 4,5) (Shanmugaraj et al., 2023b).

Published barley scRNAseq transcriptome data were queried to evaluate whether the MSI-defined metabolite domains corresponded to transcriptionally defined developmental regions (Demesa-Arevalo et al., 2026). Spatial domains corresponding to meristematic, vascular, spikelet primordium, or spikelet attachment-like regions were examined for the expression of candidate genes involved in polyamine metabolism, lysophospholipid remodeling, membrane lipid regulation, and auxin transport (Demesa-Arevalo et al., 2026). The gene IDs of the candidate genes evaluated are listed in Supplemental Table 2, 6. Spatial expression patterns were used as independent support for anatomical domain assignments.

## Statistical analysis

Bar plots, heatmaps, and statistical analyses were conducted in R version 4.5.3.

## Data availability

The MALDI-MSI datasets generated in this study will be deposited in a repository under accession number [XXXX]. Processed ion features, intensity tables, metabolite annotation tables, candidate gene lists, and scripts used for figure generation will be available at ‘XXXX’ upon publication. The RNA-seq datasets used for transcriptome integration were obtained from the European Nucleotide Archive accession numbers PRJEB51523 and PRJEB51366. Images of published imputed single-cell RNA sequencing data of barley inflorescence at W3.5 were obtained from (https://www.plabipd.de/projects/hannah_demo/Barvista.html). The gene IDs for all candidate genes analyzed in this study are provided in Supplemental Table 2-6.

## Acknowledgments

We thank C. Trautewig, K. Wolf, A. Wolf, A. Püschel, for excellent technical support; E. Geyer and the greenhouse team for plant care; L. Borisjuk and the Assimilate Allocation and NMR group at IPK for access to cryosectioning facilities; and members of the IPK Plant architecture group for discussions. This work was supported by the European Research Council (ERC) grant entitled “LUSH SPIKE” (ERC-2015-CoG, agreement 681686), the European Fund for Regional Development (EFRE), and the State of Saxony-Anhalt within the ALIVE project (grant ZS/2018/09/94616), the HEISENBERG Program of the German Research Foundation (DFG, grant Nos. SCHN 768/8-1 and SCHN 768/15-1) and the IPK core budget. Model figure was created using BioRender.com

## Author contributions

N.S., H.-P.M., and T.S. conceived the study. N.S. performed spike sampling, MALDI-MSI experiments, data analysis, hypothesis generation, and manuscript drafting. Y.H. and T.R. characterized the *hvcmf4* developmental framework, and Y.H. contributed to the *hvcmf4* transcriptome interpretation. N.S., A.G.-H., J.C. D’A. and H.-P.M. contributed to metabolite analysis, MSI methodology, validation and interpretation. T.S. and H.-P.M. supervised the study and provided funding. N.S. and T.S. wrote the manuscript with input from all authors.

## Declaration of generative AI and AI-assisted technology in writing process

Language editing and manuscript organization were assisted using ChatGPT; all scientific interpretations, and final text were reviewed and approved by the authors.

## Supplemental Data

**Supplemental Figure 1.**
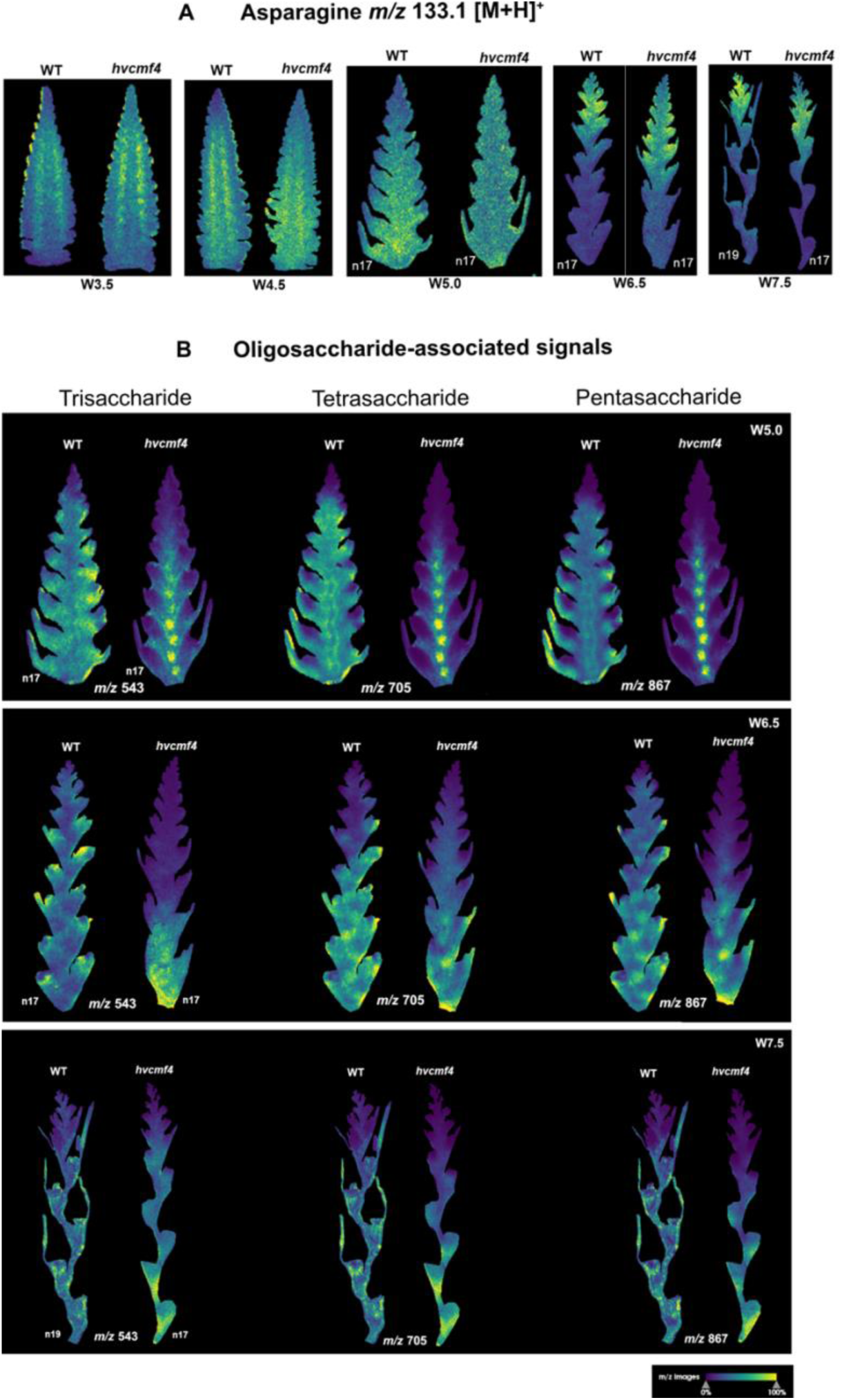
Asparagine- and oligosaccharide-associated signals distinguish degeneration-prone and viable spike regions. (A) MALDI-MSI distribution of the asparagine-associated ion *m/z* 133.1 [M+H]+ in wild-type Bowman and *hvcmf4* spikes across development. The signal preferentially accumulates in apical degeneration-prone tissues and expands over a broader apical domain in *hvcmf4*, consistent with an accelerated nitrogen-metabolic transition associated with premature PTD. (B) MALDI-MSI distribution of higher-order oligosaccharide-associated ions, including *m/z* 543.1, 705.2 and 867.3 [M+K]+, in wild-type Bowman and *hvcmf4* spikes. These signals are mainly detected during the spike growth phase and are preferentially retained in viable spikelet regions, whereas they are reduced in degenerating apical nodes, particularly in *hvcmf4*. Ion images are shown after TIC normalization, and color scales indicate relative normalized ion intensity.

**Supplemental Figure 2.**
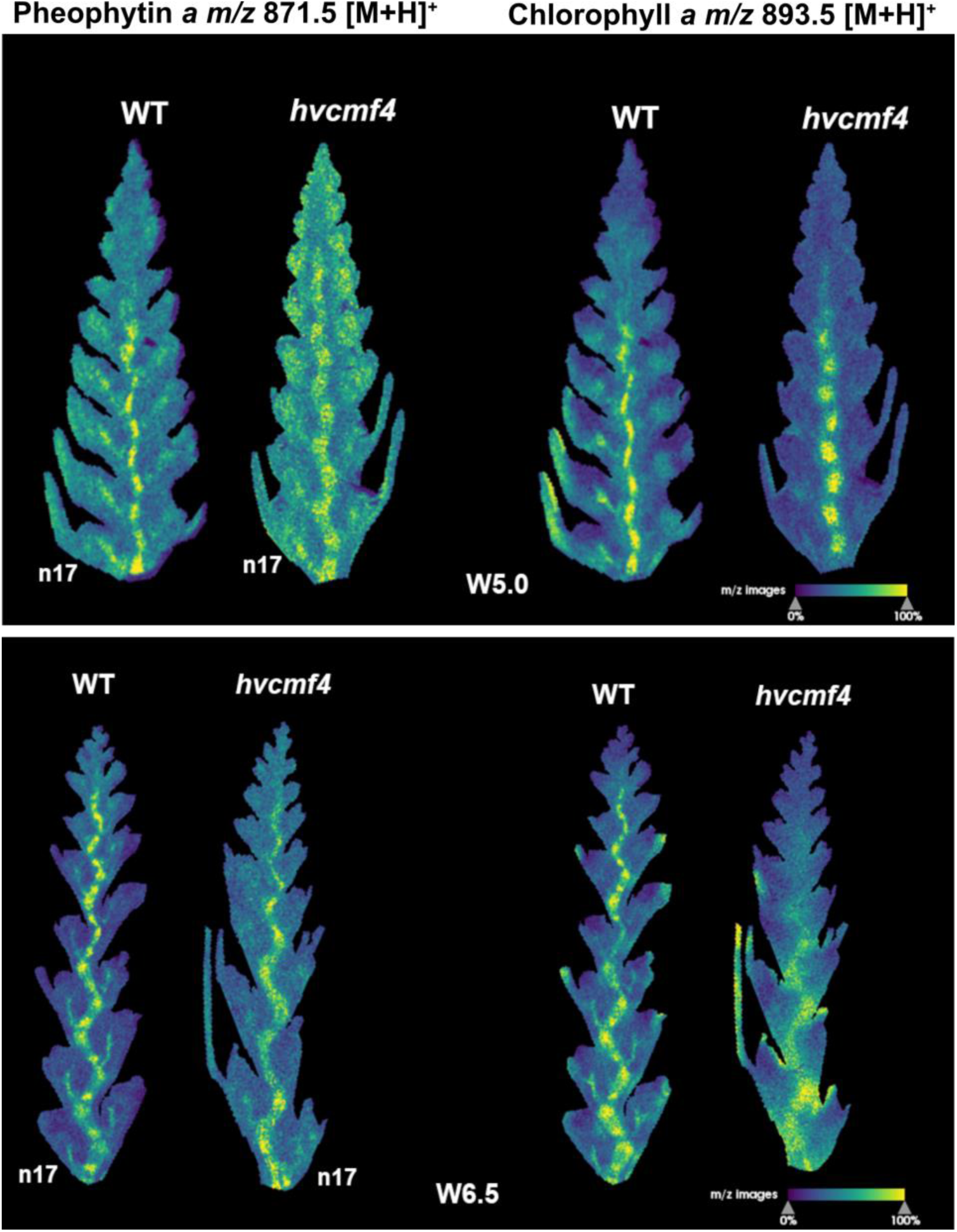
Chlorophyll- and pheophytin-associated signals reveal spatial greening patterns during barley spike development. MALDI-MSI distribution of chlorophyll-associated *m/z* 893.5 [M+H]+ and pheophytin-associated *m/z* 871.5 [M+H]+ signals in wild-type Bowman and *hvcmf4* spikes at W5.0 and W6.5. In wild-type Bowman, chlorophyll-associated signals show an acropetal pattern, consistent with progressive greening of the rachis and spikelet-associated tissues during spike growth. In *hvcmf4*, chlorophyll- and pheophytin-associated signals are reduced in spikelet-associated and apical tissues at later stages, consistent with impaired greening and premature degeneration. Ion images are shown after TIC normalization, and color scales indicate relative normalized ion intensity.

**Supplemental Figure 3.**
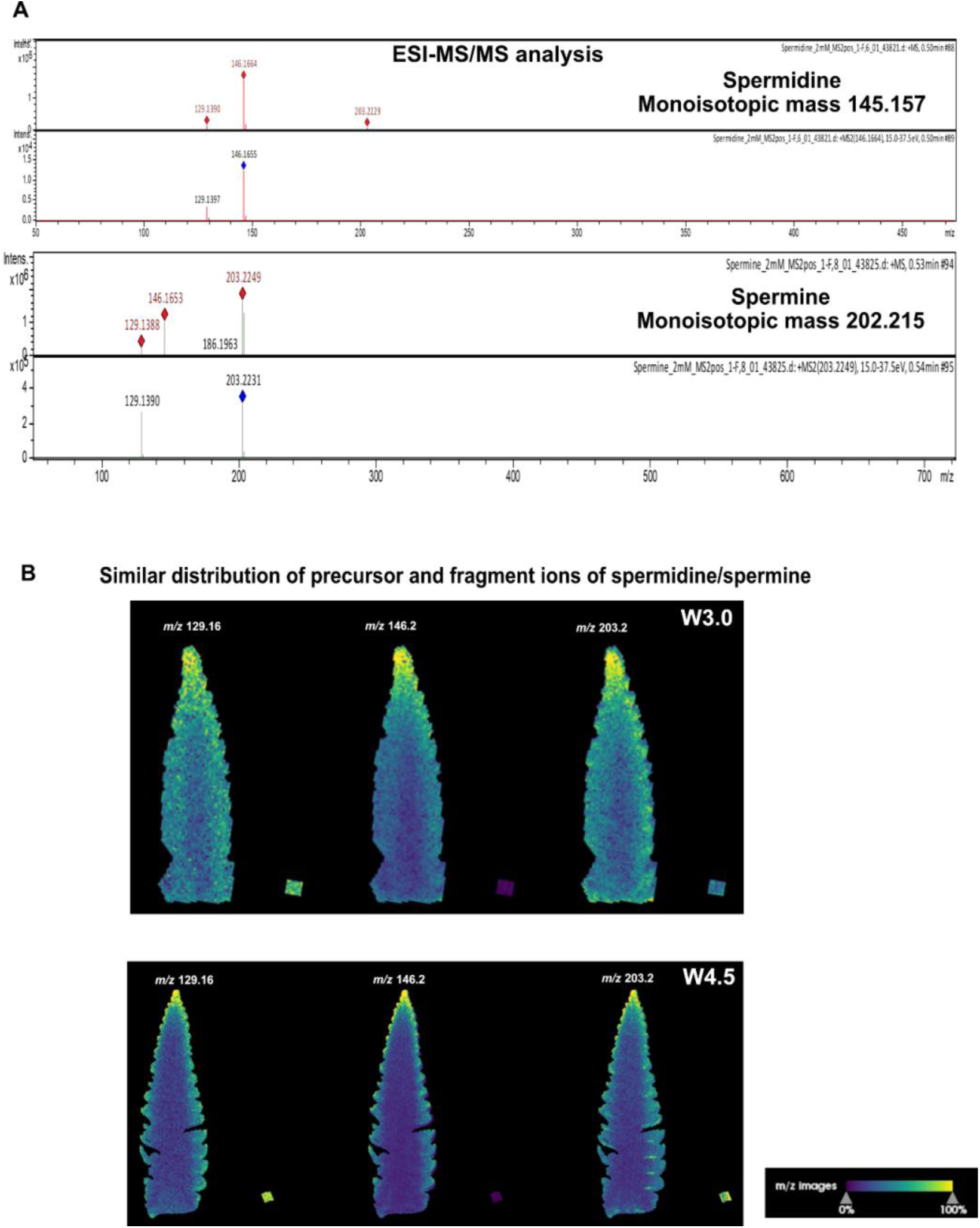
Validation and spatial distribution of spermidine- and spermine-associated ions. **(A)** ESI-MS/MS spectra of spermidine and spermine standards. Spermidine produced a dominant *m/z* 146.1 precursor ion and a related *m/z* 129.1 fragment, consistent with ammonia loss. Spermine produced a dominant *m/z* 203.2 precursor ion and related fragments, including ions overlapping the spermidine-associated mass range. These fragmentation patterns support assignment of m/z 146.1 as a spermidine-associated ion and *m/z* 203.2 as a spermine-associated ion. **(B)** MALDI-MSI ion images showing similar spatial distributions of *m/z* 129.1, *m/z* 146.2 and *m/z* 203.2 in representative W3.0 and W4.5 wild-type Bowman spike sections. The co-localization of precursor- and fragment-associated ions in meristematic apical regions supports interpretation of these features as polyamine-associated MSI signals. Ion images are shown after TIC normalization, and color scales indicate relative normalized ion intensity.

**Supplemental Figure 4.**
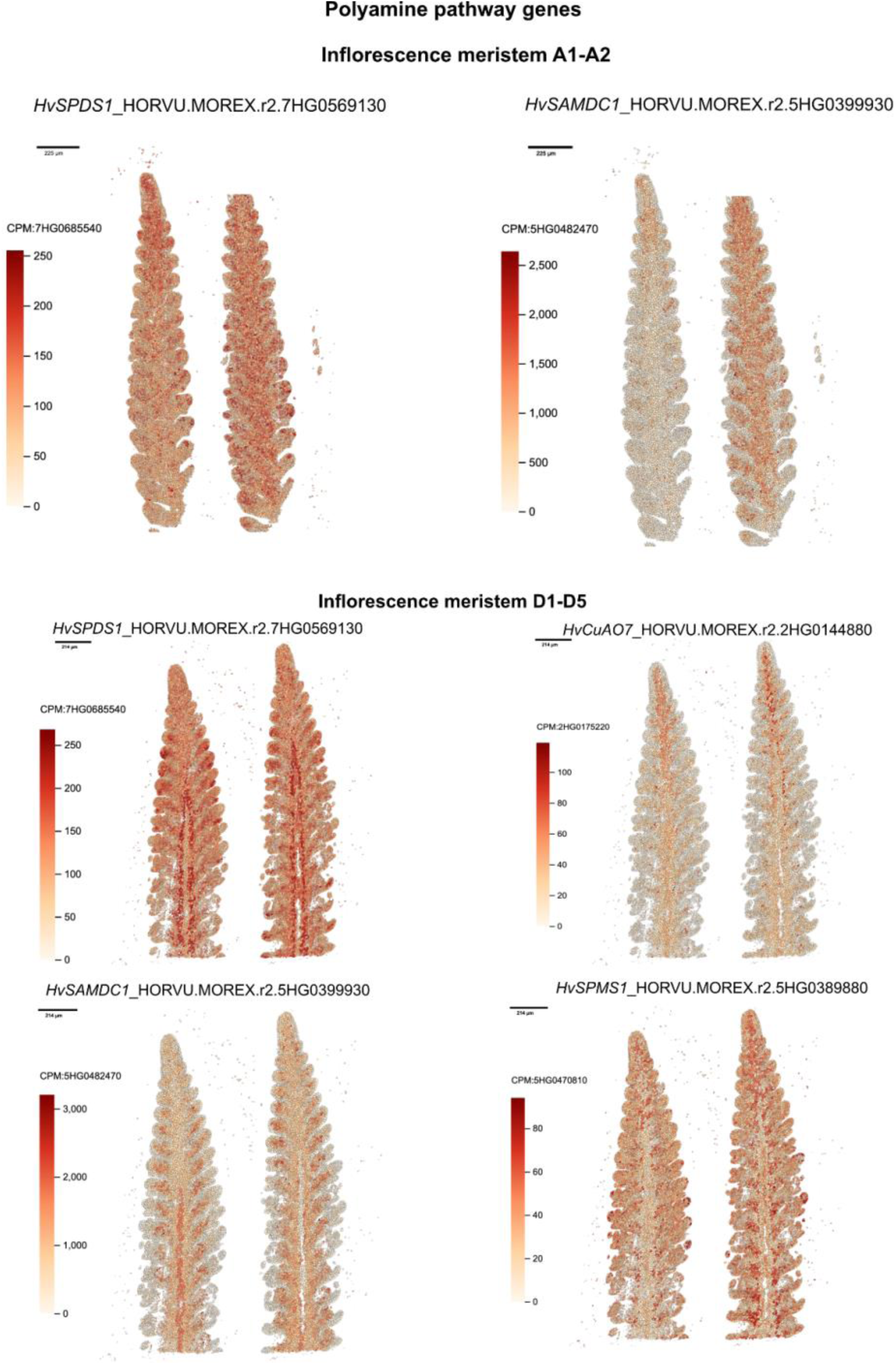
ScRNAseq and spatial transcriptome support for a meristem-associated polyamine biosynthesis domain. Published barley W3.5 imputed scRNAseq transcriptome data showing expression of candidate *HvSAMDC*, *HvSPDS*, *HvSPMS, HVCuAO* genes in meristem-associated domains, including inflorescence and spikelet meristem regions. These patterns support the interpretation that the spermidine-associated MSI signal marks a local polyamine-rich state in reproductive meristems.

**Supplemental Figure 5.**
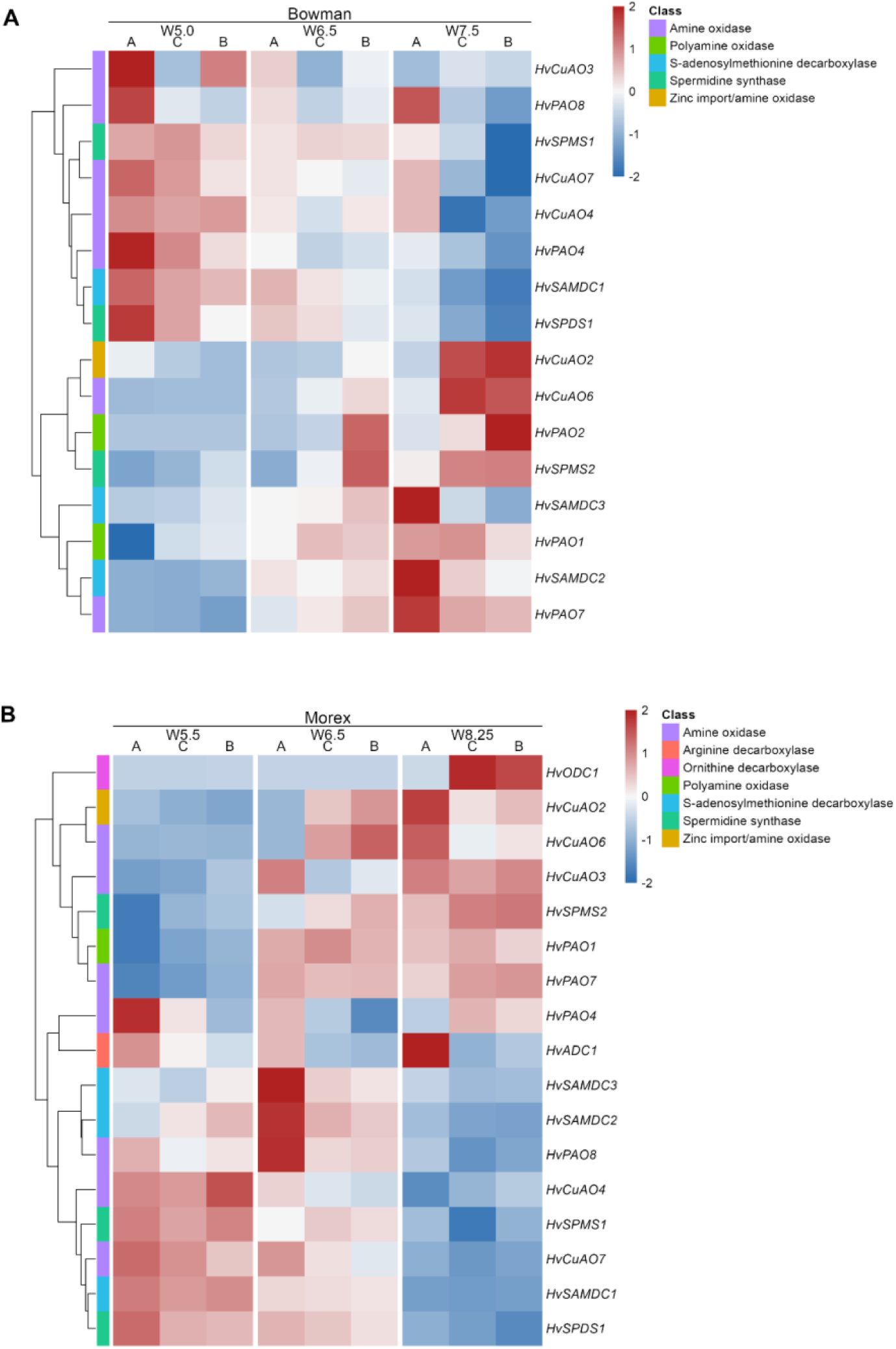
Expression patterns of candidate genes involved in polyamine biosynthesis, maintenance and oxidative turnover in previously published barley spike transcriptome datasets. **(A)** In wild-type Bowman, polyamine biosynthetic or maintenance genes, including *HvSAMDC1*, *HvSPDS1* and *HvSPMS1*, are retained in earlier active spike tissues, whereas selected *PAO*/*CuAO* genes, including *HvPAO7* and related oxidative-turnover candidates, increase in apical or degeneration-associated tissues at later PTD stages. **(B)** A comparable stage- and position-dependent remodeling of polyamine pathway genes is observed in the six-row cultivar Morex, although the timing and individual oxidative-turnover genes differ. These patterns indicate that regular developmental PTD is associated with a late transition from polyamine biosynthesis or maintenance toward polyamine oxidation and remodeling.

**Supplemental Figure 6.**
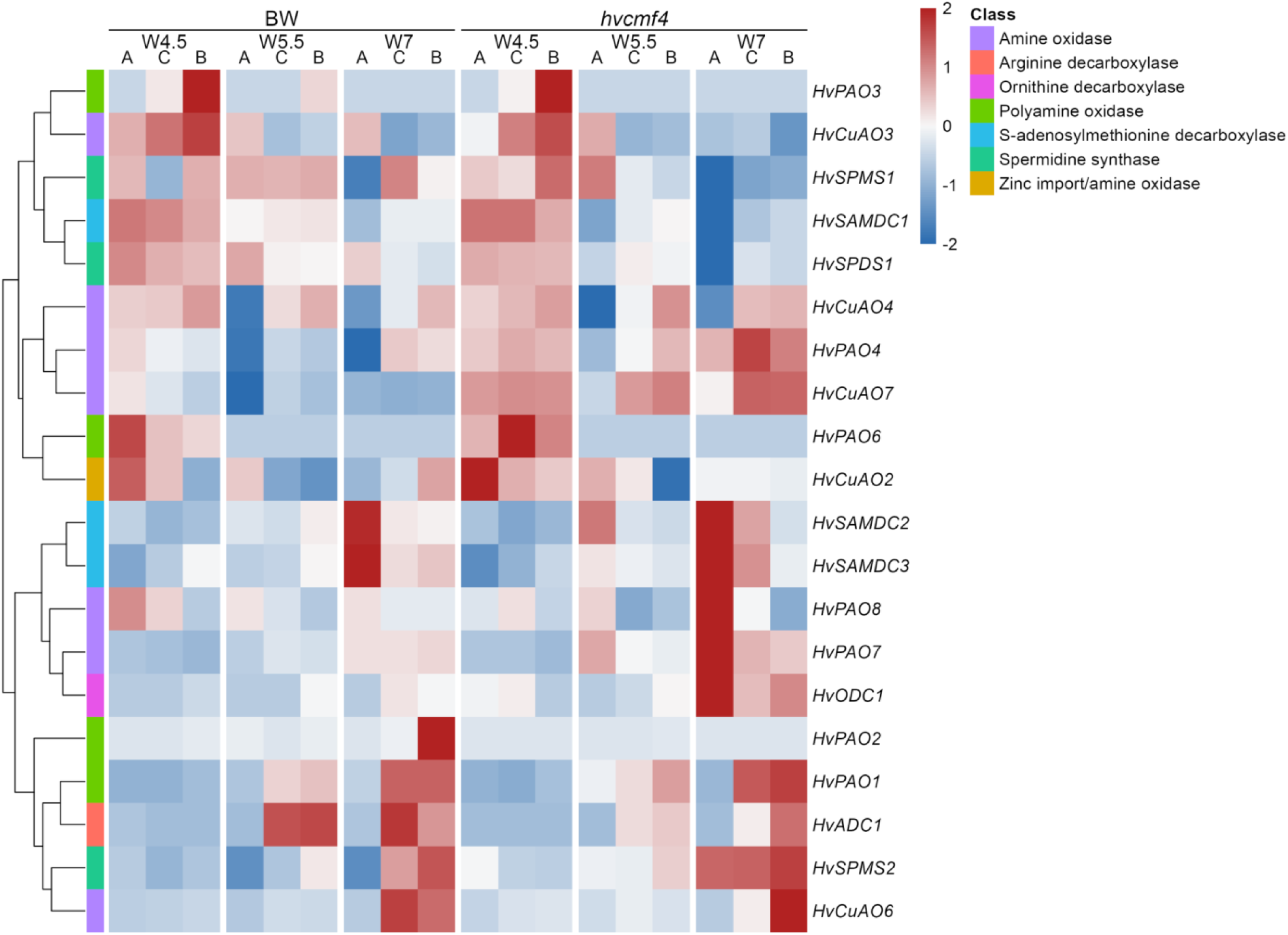
Apical polyamine pathway reprogramming in wild-type Bowman and *hvcmf4*. Published RNA-seq data showing expression of candidate polyamine biosynthesis/maintenance genes and catabolic genes across apical, central and basal spike regions in wild-type Bowman and *hvcmf4*. The strongest genotype-dependent reprogramming occurs in apical tissues, where *hvcmf4* shows reduced biosynthetic or maintenance gene expression and increased catabolic gene expression, consistent with premature activation of polyamine turnover during extended PTD.

**Supplemental Figure 7.**
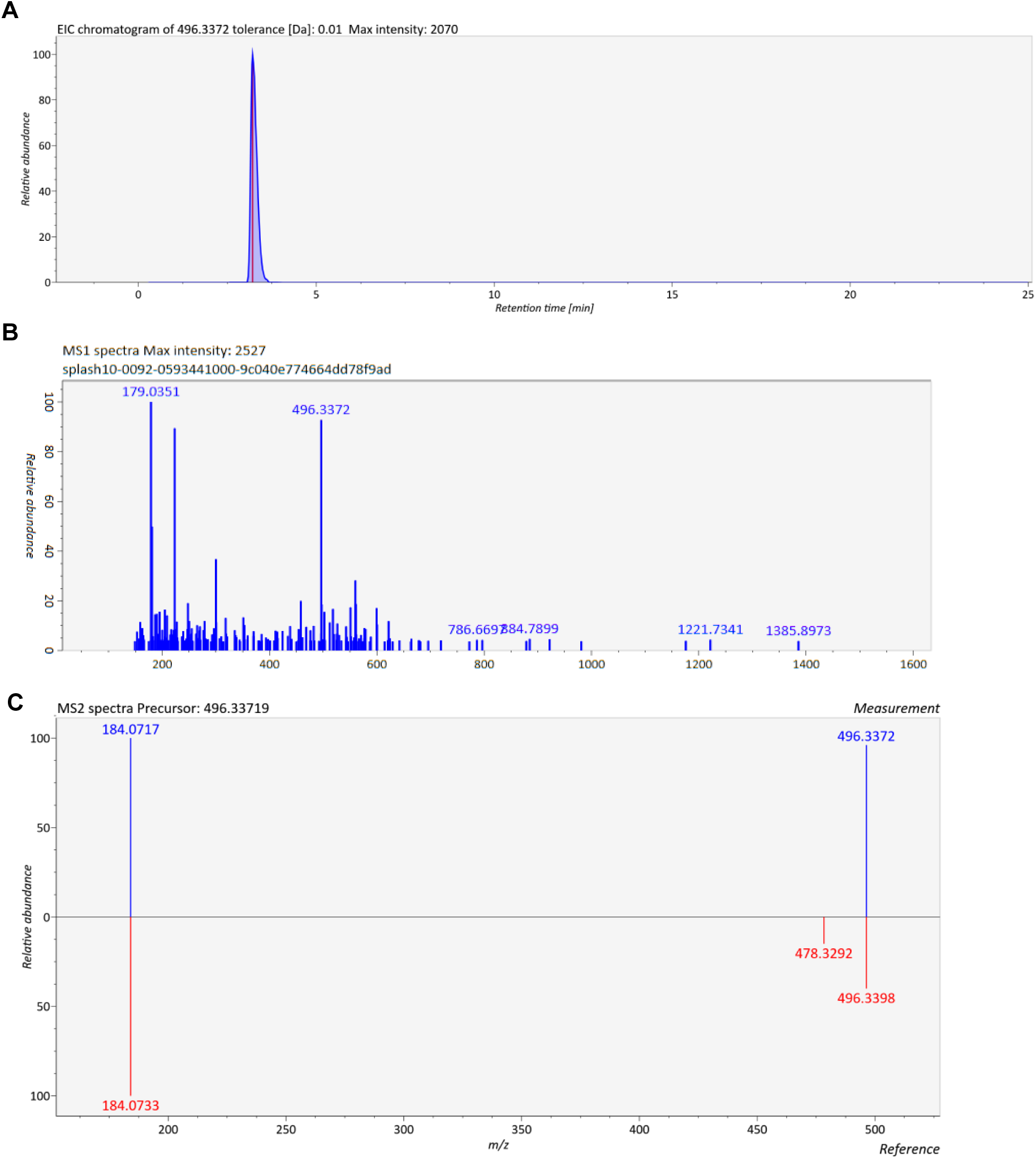
Targeted LC-ESI-MS validation of LPC-associated signals in barley spikes. (A) Extracted Ion Chromatogram (EIC) of LC-ESI-MS lipid metabolic feature *m/z* 496.33 corresponding to LPC 16:0 [M+H]^+^. (B) MS1 spectra acquired at the retention time of 3.21 min including LPC 16:0 [M+H]^+^. (C) LC-ESI-MS/MS fragmentation of the precursor *m/z* 496.33, leading to the characteristic phosphocholine group [C₅H₁₅NO₄P]⁺ *m/z* 184.07 (blue), confirmed by spectral MS matching with a reference LPC 16:0 spectra (red).

**Supplemental Figure 8.**
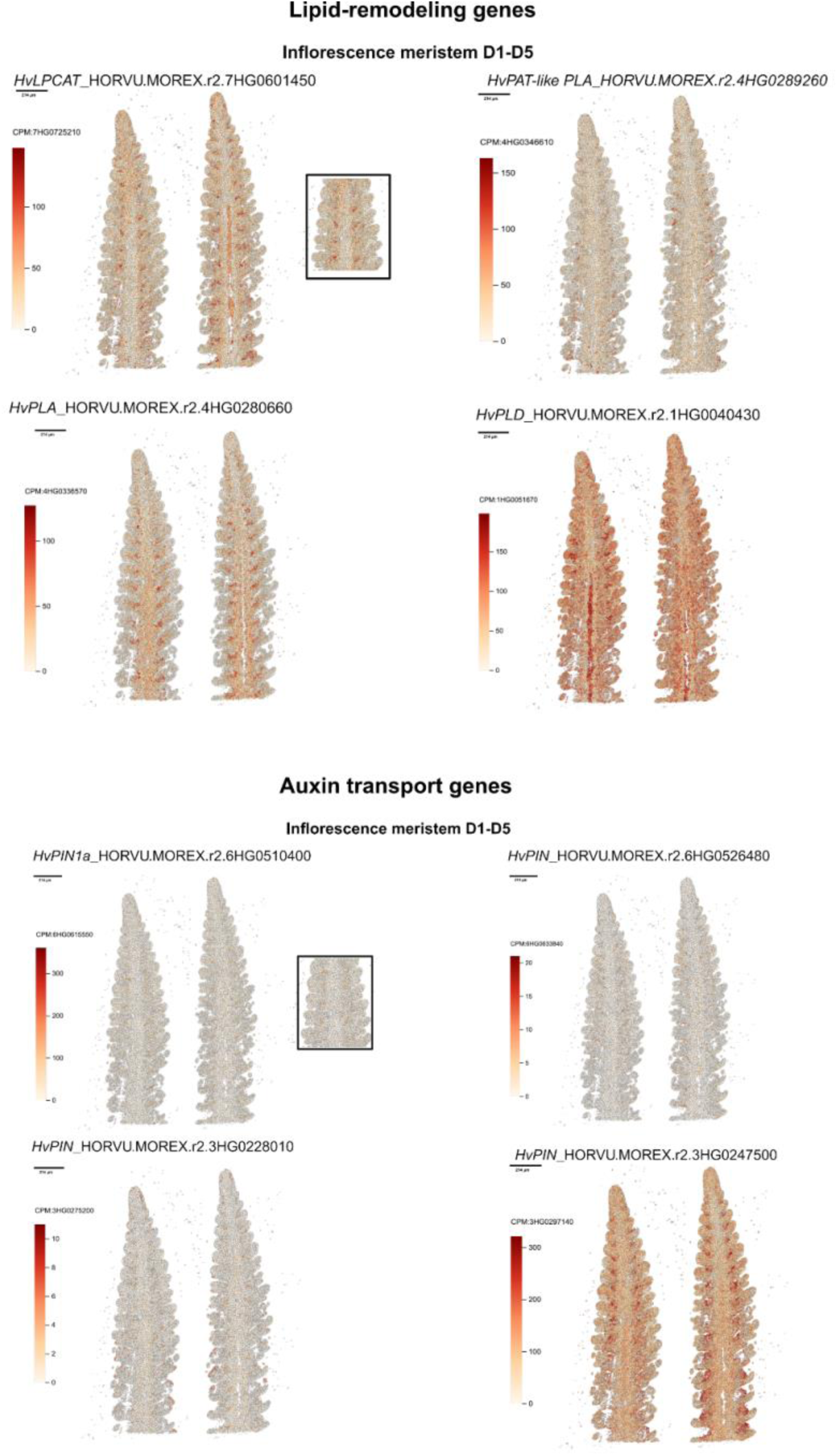
ScRNAseq and spatial transcriptome support for the LPC-associated vascular lipid-remodeling domain. Published barley W3.5 scRNAseq and imputed spatial transcriptome data were used to examine whether candidate lipid-remodeling and auxin transport genes occupy domains consistent with the LPC-associated MSI pattern. Representative candidate genes *PLA*, *LPCAT*, *PLD* and *PAT-like PLA*, together with *PIN* genes, show expression in spatial domains corresponding to the spikelet meristem base, spikelet attachment-like regions, and adjacent vascular/provascular tissues. The overlap or close adjacency of lipid-remodeling and auxin transport-associated expression domains provides independent transcriptomic support for the interpretation that the LPC-associated MSI signal marks a vascular-associated lipid-remodeling domain linked to auxin-guided supply-zone formation.

